# A *trans*-translation inhibitor is potentiated by zinc and kills *Mycobacterium tuberculosis* and non-tuberculous mycobacteria

**DOI:** 10.1101/2024.11.02.621434

**Authors:** Akanksha Varshney, Ziyi Jia, Michael D. Howe, Kenneth C. Keiler, Anthony D. Baughn

**Author notes:** These authors contributed equally (Akanksha Varshney, Ziyi Jia).

## Abstract

*Mycobacterium tuberculosis* poses a serious challenge for human health, and new antibiotics with novel targets are needed. Here we demonstrate that an acylaminooxadiazole, MBX-4132, specifically inhibits the *trans*-translation ribosome rescue pathway to kill *M. tuberculosis*. Our data demonstrate that MBX-4132 is bactericidal against multiple pathogenic mycobacterial species and kills *M. tuberculosis* in macrophages. We also show that acylaminooxadiazole activity is antagonized by iron but is potentiated by zinc. Our transcriptomic data reveals dysregulation of multiple metal homeostasis pathways after exposure to MBX-4132. Furthermore, we see differential expression of genes related to zinc sensing and efflux when *trans*-translation is inhibited. Taken together, these data suggest that there is a link between disturbing intracellular metal levels and acylaminooxadiazole-mediated inhibition of *trans*-translation. These findings provide an important proof-of-concept that *trans*-translation is a promising antitubercular drug target.

## INTRODUCTION

Infections caused by *Mycobacterium tuberculosis* have killed 30 million people in the 21^st^ century^1^. Despite an effective treatment regimen, over 1.6 million people die annually from tuberculosis (TB)^1^. Long treatment times, adverse drug reactions, and the increasing prevalence of multidrug-resistant and extensively drug-resistant strains have produced an urgent need for new antibiotics^2,3^. Drugs with new targets and novel mechanisms of action are particularly desirable to evade existing resistance mechanisms and reduce TB treatment times.

The *trans*-translation ribosome rescue pathway is a potential antibacterial target because it is essential in many pathogenic bacterial species and absent in humans. The physiological role of *trans*-translation is to rescue ribosomes that have stalled at the 3′ end of mRNAs that lack a stop codon. *trans*-Translation rescues these “non-stop” ribosomes using a specialized RNA molecule, transfer messenger RNA (tmRNA), encoded by *ssr*, and small protein B (SmpB), encoded by *smpB*^4^. Mimicking tRNA, the tmRNA-SmpB complex enters the A site of a non-stop ribosome to accept the nascent polypeptide. When tmRNA-SmpB is translocated to the P site, a specialized reading frame within tmRNA is inserted in the mRNA channel. Translation resumes using tmRNA as the message and terminates on the stop codon at the end of this reading frame, releasing the ribosome and the nascent polypeptide that now has the tmRNA-encoded sequence, known as the SsrA tag, at its C terminus^4,5^. Multiple proteases recognize the SsrA tag, resulting in rapid proteolysis of the tagged protein^6^. *trans*-Translation is an anti-tubercular drug target because it is essential for viability of growing and non-growing drug-tolerant populations of *M. tuberculosis*^6^. In addition, *trans*-translation is not targeted by any existing anti-tubercular drugs, making it unlikely that circulating clinical isolates will harbor resistance to new inhibitors.

A high throughput screen identified the acylaminooxadiazole KKL-35 as an inhibitor of *trans*-translation *in vivo* and *in vitro*^7^. However, KKL-35 was unsuitable for use in animals because its amide bond was rapidly hydrolyzed^6,8^. Optimization of pharmacokinetic and toxicity properties resulted in MBX-4132, which exhibited excellent stability in both murine liver microsomes and murine serum as well as excellent Caco-2 permeability^8^. MBX-4132 specifically inhibited *trans*-translation in *E. coli* but did not inhibit normal translation^8^. Single-particle cryogenic-electron microscopy (cryo-EM) studies using *E. coli* ribosomes revealed that the acylaminooxadiazoles bind near the peptidyl transfer center^8^. In addition, MBX-4132 could clear multi-drug resistant *Neisseria gonorrhoeae* from infected mice after a single oral dose, demonstrating that inhibition of *trans*-translation is a viable antibiotic target for treating drug-resistant infections^8^.

Here, we report that MBX-4132 specifically inhibits *M. tuberculosis trans*-translation *in vitro* and is bactericidal against *M. tuberculosis*, *Mycobacterium avium,* and *Mycobacterium abscessus.* Our studies reveal that iron has antagonistic effects on the activity of acylaminooxadiazoles but zinc can overcome these effects and potentiate the antibacterial activity. We have also examined the transcriptional responses induced by MBX-4132 treatment, characterized MBX-4132 activity against a tmRNA knockdown mutant strain, and identified transposon insertion mutants and a deletion strain with altered susceptibility, which sheds light on the ability of MBX-4132 to both impact metal homeostasis and interact with ribosomes in *M. tuberculosis*. Further, we characterize the antitubercular activity of MBX-4132 in a macrophage model of infection. Taken together, these studies highlight acylaminooxidiazoles as potent on-target lead inhibitors of *trans*-translation for the discovery of new antitubercular agents.

## RESULTS

### MBX-4132 inhibits *M. tuberculosis trans*-translation *in vitro*

To determine if acylaminooxadiazole compounds can inhibit *trans*-translation on *M. tuberculosis* ribosomes, genes encoding ten translation factors and SmpB from *M. tuberculosis* were cloned and the proteins were purified from over-expressing strains of *E. coli*. Ribosomes were purified from *M. tuberculosis*, and *M. tuberculosis* tmRNA was transcribed *in vitro* and purified. These components were combined with tRNAs, aminoacyl-tRNA synthetases, methionyl-tRNA formyltransferase, and nucleoside diphosphate kinase purified from *E. coli,* and purchased T7 RNA polymerase, nucleoside triphosphates, amino acids, and salts, to produce a reaction mixture capable of *trans*-translation. The reaction mixture was incubated with [^35^S]-methionine and DNA encoding dihydrofolate reductase (DHFR) with or without an in-frame stop codon, and newly synthesized protein was observed by SDS-PAGE followed by phosphorimaging (Fig. S1). When the template included an in-frame stop codon a single protein product was observed. When the template lacked a stop codon a second, larger band was observed, consistent with the addition of the tmRNA-encoded peptide tag to DHFR. As expected, the abundance of the larger band decreased substantially when tmRNA-SmpB was omitted from the reaction and when a DNA oligonucleotide complementary to the *M. tuberculosis* tmRNA coding sequence was included, confirming that the larger band is the result of *trans*-translation.

The amount of *trans*-translation was decreased by the inclusion of KKL-35 or MBX-4132 in the reaction (Figs. 1 & S2). Dose-response experiments showed that MBX-4132 inhibited *trans*-translation with an IC_50_ = 13 ± 1 µM (Fig. 1). MBX-4132 did not inhibit normal translation (Fig. 1C). These results demonstrate that MBX-4132 specifically inhibits *trans*-translation and not translation by *M. tuberculosis* factors, similar to its activity in reactions with *E. coli* factors.

**Fig 1.**
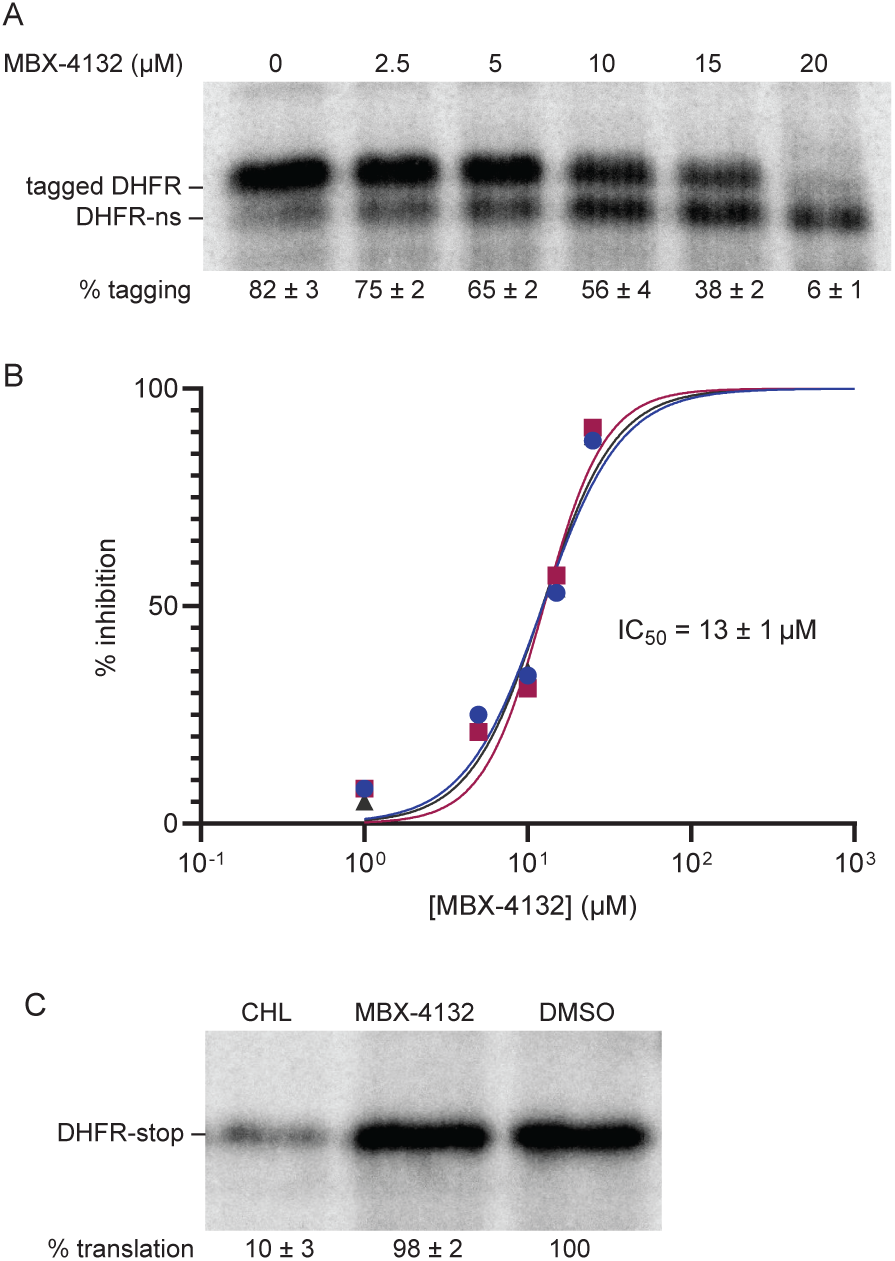
MBX-4132 inhibits *M. tuberculosis trans*-translation *in vitro.* A) A gene encoding DHFR without a stop codon was expressed in the presence of *M. tuberculosis* tmRNA-SmpB and varying concentrations of MBX-4132. Synthesized protein was detected by incorporation of ^35^S-methionine followed by SDS-PAGE and phosphorimaging. Bands corresponding to tagged and untagged DHFR are indicated, and the average percentage of DHFR protein found in the tagged band for three repeats is shown with the standard deviation. B) Data from gels as in (A) were plotted and fit with a sigmoidal function to determine the IC_50_. C) *in vitro* translation was assayed from the expression of a gene encoding DHFR with a stop codon in the presence of DMSO, 20 μM chloramphenicol (CHL), or 20 μM MBX-4132, and a representative experiment is shown. The percentage of DHFR with respect to the amount in the DMSO-treated control is shown as the average from two independent repeats with the standard deviation.

### MBX-4132 competes with mycobacterial tmRNA-SmpB

To test whether MBX-4132 competes with tmRNA-SmpB during inhibition, we added excess *M. tuberculosis* tmRNA-SmpB to the *in vitro trans*-translation reactions. In reactions with 10 µM MBX-4132, increasing the tmRNA-SmpB concentration from 2.75 µM to 8.2 µM decreased inhibition from 63% to 7%. When the MBX-4132 concentration was increased to 20 µM in reactions with 8.2 µM tmRNA-SmpB, inhibition increased to 85% (Fig. 2). These data indicate that MBX-4132 competes with tmRNA-SmpB activity on the *M. tuberculosis* ribosome.

**Fig 2.**
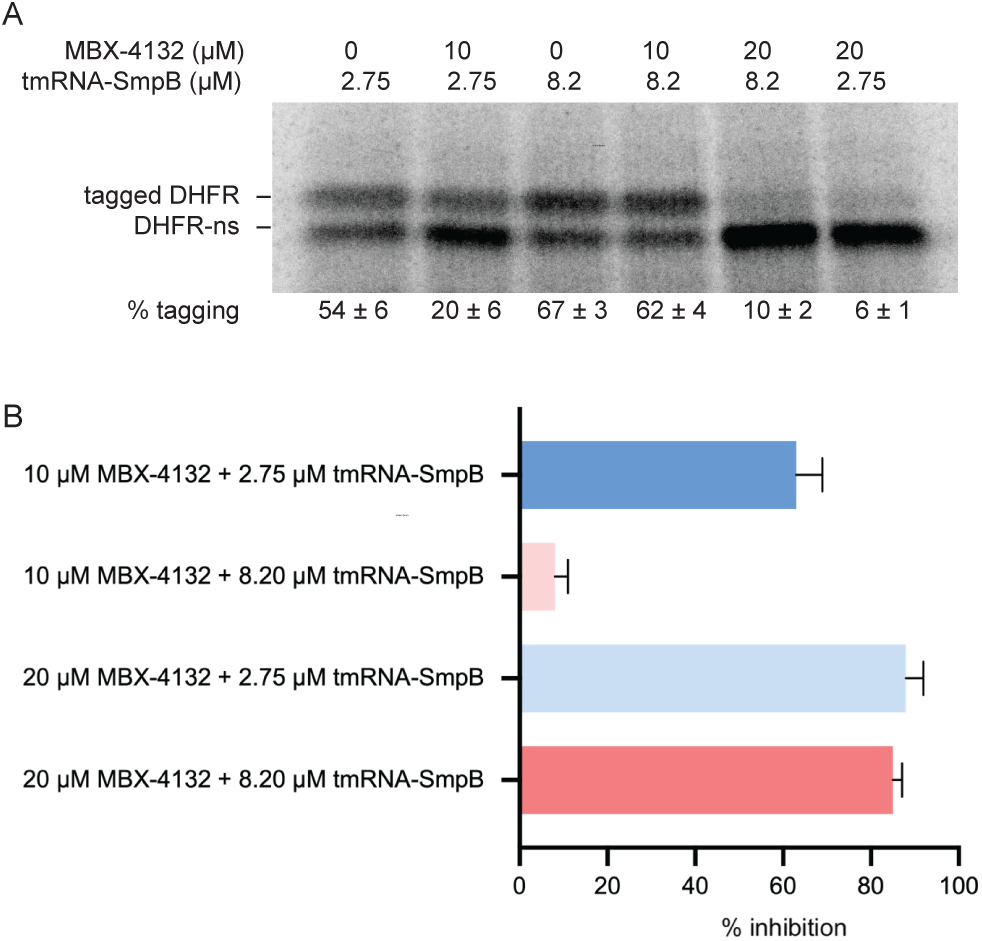
tmRNA-SmpB competes with MBX-4132 *in vitro*. A) *in vitro trans*-translation assays as in Figure 1 containing different concentrations of tmRNA-SmpB and MBX-4132. B) Reactions treated with 8.2 µM tmRNA-SmpB suppressed the inhibition of *trans*-translation by 10 µM MBX-4132. The inhibition was rescued by 20 µM MBX-4132. Data from at least two experiments are shown as the average with error bars indicating the standard deviation.

### MBX-4132 has anti-mycobacterial activity

To assess the anti-mycobacterial activity of KKL-35 and MBX-4132, we performed broth dilution and plating assays (Table 1). MBX-4132 has a similar potency to KKL-35 against many bacterial pathogens^8^, but neither KKL-35 nor MBX-4132 inhibited the growth of *M. tuberculosis* H37Rv Δ*RD1* Δ*panCD* in broth dilution assays in Middlebrook 7H9 medium. Proteomic profiling has demonstrated that *trans*-translation inhibitors can disturb metal homeostasis in *E. coli* and *Bacillus subtilis*^9^, so we examined the impact of metals on oxadiazole activity using a defined minimal medium (Mtb Minimal Medium, MM)^10^, made without addition of ferric ammonium citrate. In this medium, hereafter referred to as low-iron Mtb Minimal Medium (LIMM), KKL-35 was bactericidal against *M. tuberculosis* (MBC = 0.4 µg/mL), *M. avium* (MBC = 3.1 µg/mL) and *M. abscessus* (MBC = 6.4 µg/mL) (Table 1). Likewise, MBX-4132 was bactericidal against *M. tuberculosis* (MBC = 1.6 µg/mL), *M. avium* (MBC = 2.1 µg/ml), and *M. abscessus* (MBC = 3.3 µg/mL) in LIMM (Table 1). This growth inhibition in the absence of added iron suggests that acylaminooxadiazoles kill mycobacteria by inhibiting *trans*-translation but are counteracted by the presence of iron.

**Table 1.** Minimum inhibitory and minimum bactericidal concentrations of *trans-*translation inhibitors for mycobacterial species.

| Compound | *M. tuberculosis* |  | *M. avium* |  | *M. abscessus* |  | *E. coli* ΔtolC |  |
| --- | --- | --- | --- | --- | --- | --- | --- | --- |
|  | MIC<sup>a</sup> | MBC<sup>b</sup> | MIC | MBC | MIC | MBC | MIC | MBC |
| KKL-35 | 0.4<br>(1.25) | 0.4<br>(1.25) | 3.1<br>(9.7) | 3.1<br>(9.7) | 6.3<br>(19.8) | 6.3<br>(19.8) | 0.4<br>(1.25) | 0.4<br>(1.25) |
| MBX-4132 | 0.8<br>(2.5) | 1.6<br>(5) | 2.1<br>(6.2) | 2.1<br>(6.2) | 3.3<br>(9.7) | 3.3<br>(9.7) | 0.8<br>(2.5) | 1.6<br>(5) |
| Rifampicin | 0.12<br>(0.14) | 0.12<br>(0.14) | 0.5<br>(0.6) | 1<br>(1.2) | 128<br>(155) | 256<br>(311) | ND<sup>c</sup> | ND |
| Azithromycin | 4<br>(5.3) | ND | 32<br>(42.4) | ND | 8<br>(10.6) | ND | ND | ND |

<sup>a</sup> MIC; µg/mL (µM) values from at least three broth microdilution assays.
<sup>b</sup> MBC; µg/mL (µM) values from at least three plating assays.
<sup>c</sup> ND: Not determined.

### Addition of zinc and removal of iron potentiate the anti-mycobacterial activity of MBX-4132

Zinc supplementation can potentiate antibacterial activity of some compounds^11–14^. To assess the effect of Zn^2+^ on the anti-mycobacterial activity of MBX-4132, we performed plating assays in 7H10 agar supplemented with ZnSO_4_. Zinc potentiated MBX-4132 against *M. tuberculosis* and *M. avium* but had much less of an impact on its activity against *M. abscessus* (Table 2). Similar results were obtained when broth microdilution assays were repeated in LIMM supplemented with ZnSO_4_ (Table 3). Results from these experiments reveal that zinc can overcome the antagonistic effects of iron and potentiate the antibacterial activity of MBX-4132.

**Table 2.**
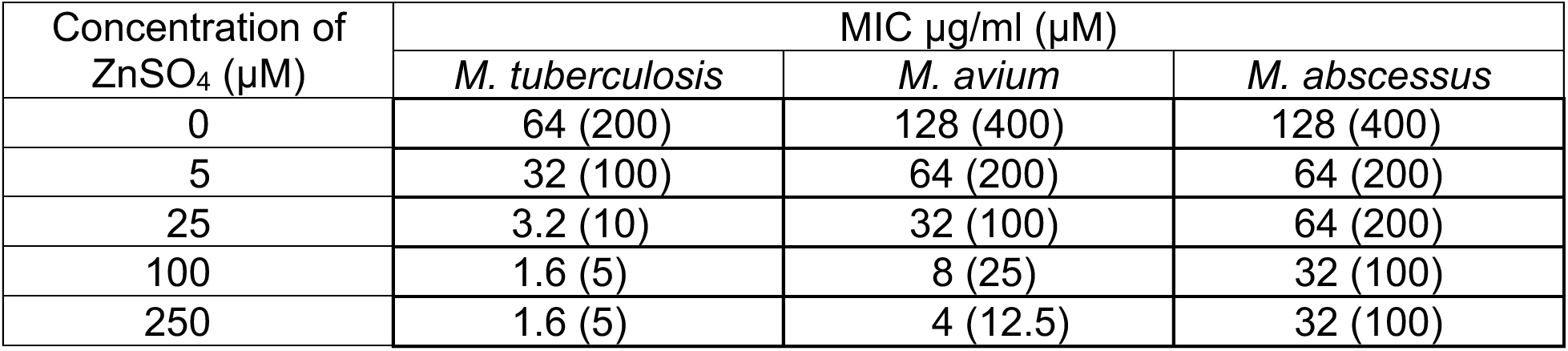
Effect of Zn^2+^ on antibacterial activity of MBX-4132 in 7H10 agar.

**Table 3.** Effect of Zn^2+^ on antibacterial activity of MBX-4132 in LIMM.

| Concentration of ZnSO<sub>4</sub> (μM) | MIC μg/ml (μM) |  |  |
| --- | --- | --- | --- |
|  | *M. tuberculosis* | *M. avium* | *M. abscessus* |
| 0 | 1.6 (5) | 2.1 (6.2) | 3.3 (10) |
| 5 | 1.6 (5) | 2.1 (6.2) | 3.3 (10) |
| 25 | 0.8 (2.5) | 1.05 (3.1) | 3.3 (10) |
| 100 | 0.4 (1.2) | 1.05 (3.1) | 3.3 (10) |
| 250 | 0.4 (1.2) | 1.05 (3.1) | 1.6 (5) |

### Iron decreases the activity of MBX-4132 *in vitro*

Growth inhibition assays demonstrated that iron decreases the activity of MBX-4132 and activity is restored by omitting iron from the culture medium. In principle, iron could have a physiological effect on the cells, for example, through alteration of cell envelope permeability. Alternatively, iron could directly interact with MBX-4132 thereby affecting the inhibition of *trans*-translation. To assess if iron affects the inhibition of *trans*-translation *in vitro*, we incubated 150 µM Fe_2_(SO_4_)_3_ with 15 µM MBX-4132 or KKL-35 and added the mixture to *in vitro trans*-translation reactions (Fig. 3). The inclusion of iron decreased the inhibition of *trans*-translation from 62% to 6% for MBX-4132, and from 36% to 4% for KKL-35 (Fig. 3). When the iron chelator TPEN was included in the incubation, inhibition of *trans*-translation was restored, demonstrating that the availability of Fe^3+^ ions was responsible for interfering with MBX-4132 and KKL-35 (Fig. 3). FeSO_4_ also counteracted inhibition, indicating that the oxidation state of the added iron is not critical (Fig. 3). Collectively, these results demonstrate that iron directly affects acylaminooxadiazole inhibition of *trans*-translation. Unlike iron, zinc had no effect on inhibition of *trans*-translation *in vitro* (Fig. S3), suggesting that potentiation of acylaminooxadiazoles by zinc is likely due to physiological effects on mycobacteria.

**Fig 3.**
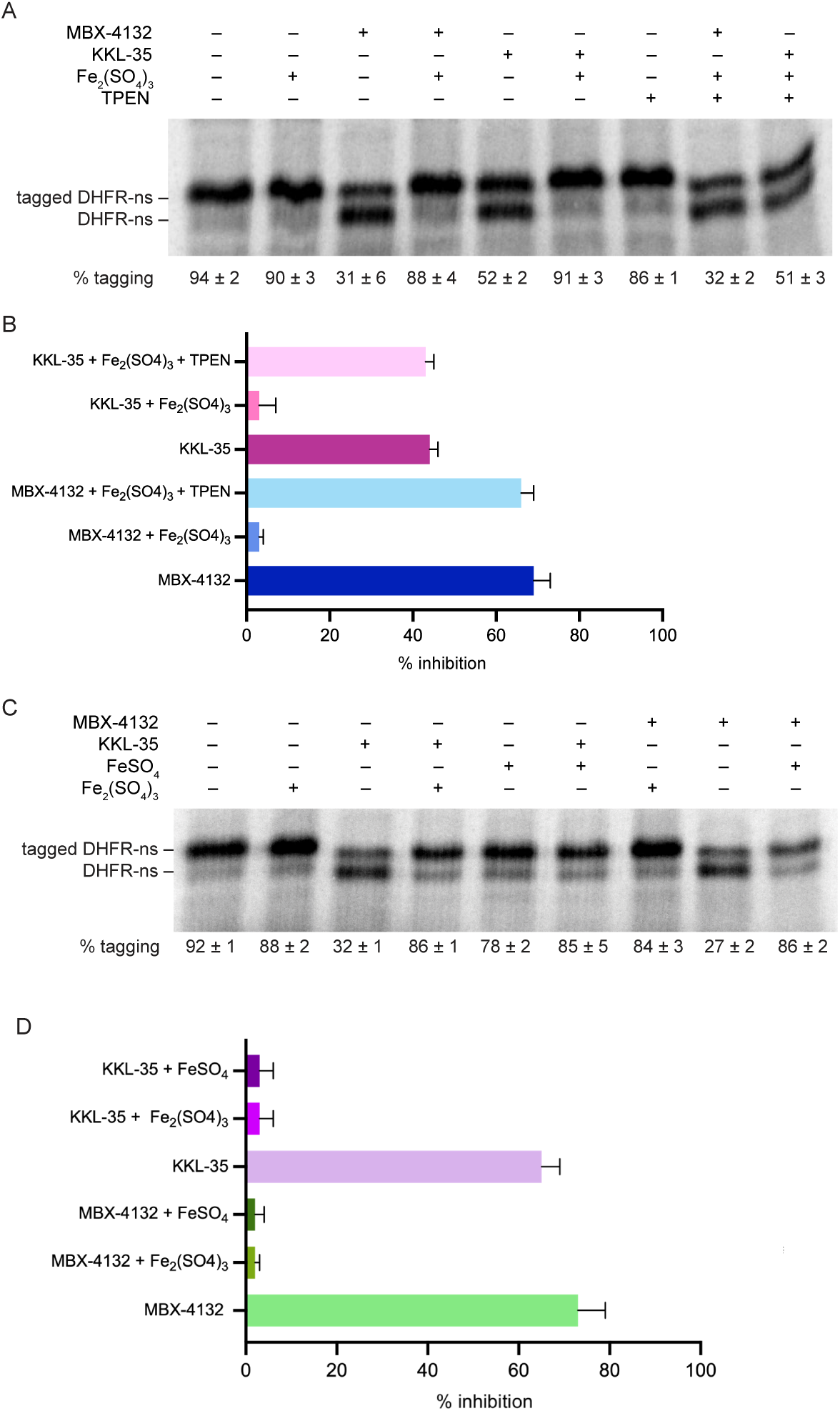
Iron inhibits the activity of MBX-4132. A) *in vitro trans*-translation assays as in Figure 1 containing 150 µM Fe_2_(SO_4_)_3_, 150 µM TPEN, 15 µM MBX-4132, or 15 µM KKL-35 as indicated. The average percentage tagging from two independent reactions is shown with the standard deviation. B) Data from gels as in (A) were plotted to show the average from two experiments with error bars indicating the standard deviation. C) *in vitro trans*-translation assays as in Figure 1 containing 150 µM Fe_2_(SO_4_)_3_, 150 µM FeSO_4_, 15 µM MBX-4132, or KKL-35 as indicated. D) Data from gels as in (C) were plotted to show the average from two experiments with error bars indicating the standard deviation.

### MBX-4132 treatment induces broad transcriptional responses

To explore the response of *M. tuberculosis* to MBX-4132 *in vivo*, we treated exponentially growing H37Rv with 1.2 µM MBX-4132 (Fig. S4A) or an equal volume dimethyl sulfoxide (DMSO) for 48 h in high zinc Mtb Minimal Medium (HZMM), prepared by supplementing MM with 3.5 μM ZnSO_4_^10^, and used RNA sequencing (RNA-seq) to compare transcript levels. Principal component analysis (PCA) showed tight clustering by biological replicates (Fig. S4B). Using a threshold of 2-fold change with adjusted *p-*value <0.05, we observed that 599 genes were upregulated (15% of annotated coding sequences) and 247 were downregulated (6% of annotated coding sequences) upon exposure to MBX-4132 (Fig. 4A, Dataset S1, Note S1).

**Fig 4.**
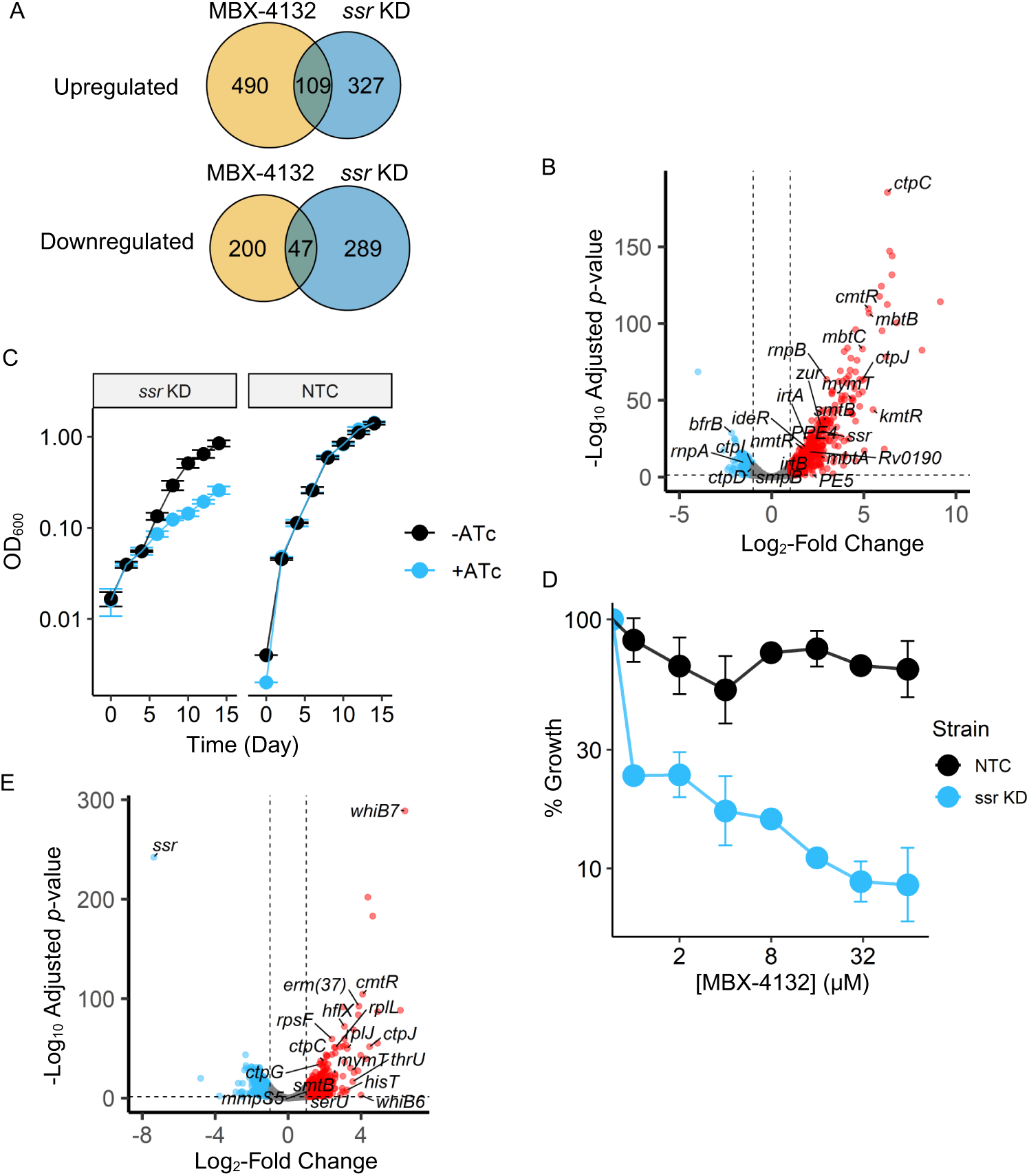
Transcriptional responses of *M. tuberculosis* H37Rv to MBX-4132 treatment and tmRNA knockdown. A) Venn diagrams showing numbers of genes significantly up- and down-regulated in the MBX-4132 study, the *ssr* KD study, or both. B) Differential expression of genes in response to MBX-4132 exposure, with vertical and horizontal dashed lines representing log_2_-fold change cutoff of ±1 and adjusted *p-*value cutoff of 0.05, respectively. C) Growth of *M. tuberculosis* H37Rv *ssr* KD and NTC strains upon induction in HZMM. Data represents the average OD_600_ of biological triplicates with error bars denoting the standard deviation. D) Inhibition of MBX-4132 against *M. tuberculosis* H37Rv *ssr* KD and NTC strains after 21 days of treatment in MM. Data represent geometric means and geometric standard deviations for 3 biological replicates. E) Differential expression of genes as a result of *ssr* KD, with dashed lines denoting cutoffs as described in (B).

tmRNA levels increased >4-fold, consistent with cells sensing a deficit in *trans*-translation (Fig. 4B & S5A). In contrast, the *smpB* transcript did not change significantly (Fig. 4B & S5A). Likewise, the RNA component of RNase P, encoded by *rnpB*^15^, was more abundant, but not *rnpA*, which encodes the protein subunit^16^ (Fig. 4B & S5A). RNase P participates in the maturation of tRNA and tmRNA^17^, so increased tmRNA and RNase P may both result from cells attempting to increase the amount of *trans*-translation.

Genes involved in metal homeostasis were also differentially regulated. *bfrB*, which is responsible for storing excess iron, was strongly repressed^18^, whereas *ideR*, the iron-sensing transcriptional regulator, and most genes associated with siderophore-mediated iron uptake (*mbtA-mbtM, mmpL5, mmpS5*, the ESX-3 operon, and *irtAB*)^19^, were significantly induced (Fig. 4B & S5A). Member genes of the RicR (*Rv0190*) regulon, a crucial mediator of copper metabolism^20^, were also upregulated (Fig. 4B & S5A).

Transcript levels of several other metal-sensing transcriptional regulators were likewise found to have increased, including *smtB* (*Rv2358*), *zur*, *kmtR, cmtR, cadI,* and *nmtR*^21^ (Fig. 4B & S5A), but their cognate regulons were not differentially expressed. In addition, several genes encoding for metal efflux pumps (*ctp* genes)^22^ were differentially expressed in a less coherent pattern. Overall, these observations suggest that both *trans*-translation and metal homeostasis were perturbed when *M. tuberculosis* was treated with MBX-4132.

To determine whether the apparent impacts on metals were connected to inhibition of *trans-*translation, we engineered a hypomorphic knockdown (KD) strain of *ssr* in *M. tuberculosis* H37Rv using CRISPRi (Clustered Regularly Interspaced Short Palindromic Repeats interference), as well as a non-targeting control (NTC) strain^23^. The KD strain grew slowly after 6 days of induction in HZMM, reaffirming the essentiality of *trans*-translation *in vivo* (Fig. 4C). Moreover, after 7 days of induction (Fig. S4C), the *ssr* KD strain showed a moderate level of susceptibility to MBX-4132 in MM (IC_90_ = 31 μM), whereas the NTC strain was resistant (IC_90_ > 250 μM) (Fig. 4D).

Next, we characterized transcriptional responses of *M. tuberculosis* to tmRNA KD after a 7-day induction period in HZMM (Fig. S4D). We again observed tight clustering by biological replicates in PCA (Fig. S4E). Using the same log_2_-fold change and adjusted *p*-value cutoffs, we found that 436 were upregulated (11% of annotated coding sequences) and 336 were downregulated (9% of annotated coding sequences) (Fig. 4A, Dataset S1 Note S1). Most notably, tmRNA levels were reduced by 165-fold, further validating the KD system (Fig. 4E). A large number of genes encoding ribosomal proteins were also significantly upregulated, indicating that *M. tuberculosis* sensed an increased need for ribogenesis due to impaired ability to rescue stalled ribosomes (Fig. 4E).

Clustering heatmap plots directly comparing both studies show that MBX-4132-treated and *ssr* KD replicates clustered closer to each other than to their respective controls (Fig. S4G), and there were 109 coding sequences induced and 47 repressed by both MBX-4132 and knocking down tmRNA (Fig. 4A, Dataset S1, Note S1), indicating that the former produced highly similar transcriptomic responses to the latter. Most prominently, induction of genes associated with transcriptional and translational pausing (*whiB6*, *whiB7, erm(37),* and *hflX*)^24–26^, was observed in both studies (Fig. 4A&E). Upregulated genes also include those that encode metal-sensing proteins (*cadI, cmtR, smtB,* and *mymT*), and efflux pumps (*ctpC, ctpG,* and *ctpJ*) (Fig. 4A&E).

On the other hand, only one iron-related gene, *mmpS5,* was found to be differentially expressed in the *ssr* KD strain (Fig. 4E). Similarly, pathways related to iron uptake and utilization were revealed to be significantly enriched upon MBX-4132 treatment but not *ssr* KD by gene ontology analyses (Fig. S5B&C). These findings indicate that some aspects of the metal homeostasis dysregulation seen after MBX-4132 treatment are likely due to inhibition of *trans*-translation, but dysregulation of the iron-responsive genes is caused by off-target effects.

### Numerous pathways impact MBX-4132 susceptibility

To identify genes that impact MBX-4132 susceptibility and resistance, we subjected a saturated *M. tuberculosis* H37Rv transposon insertion library to a subinhibitory concentration of MBX-4132 (Fig. S6) and assessed the changes in relative abundance and fitness of cells using transposon insertion sequencing (Tn-Seq). Using the resampling model of TRANSIT^27^, our data revealed 33 genes that conferred fitness gain (log_2_-fold change > 1) and 27 genes that resulted in fitness loss (log_2_-fold change < -1) and met the significance threshold (adjusted *p*-value < 0.05). The majority of genes with a known function in which transposon insertions conferred decreasing fitness during treatment with MBX-4132 have roles in translation control (Dataset S2). These include *truA*, a tRNA pseudouridine synthase that is important for translation accuracy and efficiency^28^, *lepA*, a ribosome biogenesis factor^29^, *ychF* (*Rv1112*), which regulates translation during stress^30^, *relA*, a mediator of the stringent response^31^, and *ppk1*, a polyphosphate kinase that affects numerous cellular processes during stress, including ribosome function^32,33^. Similar genes have been identified as synthetically impaired in Tn-Seq screens with *ssrA* deletion strains of other bacteria^34,35^ (Fig. 5A). Decreased fitness of these mutants during exposure to MBX-4132 is consistent with synergistic effects from inactivating multiple translation regulatory mechanisms with inhibition of *trans-*translation.

**Fig 5.**
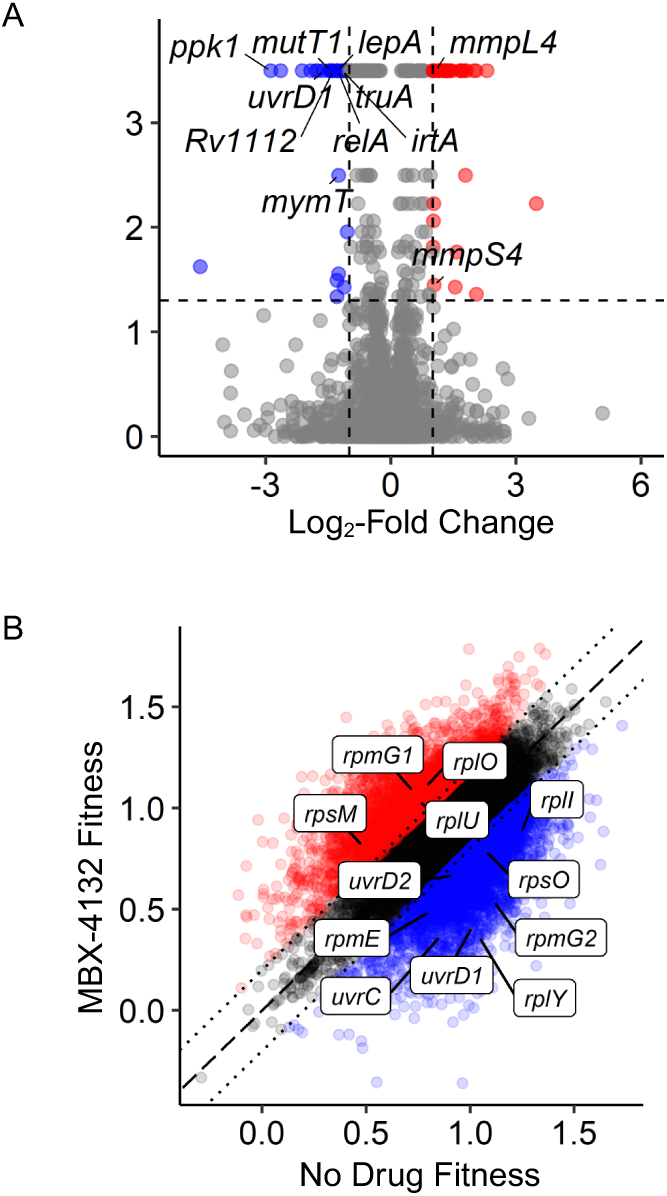
Changes of fitness of a saturated *M. tuberculosis* H37Rv transposon library after MBX-4132 treatment. A) Volcano plot showing insersions conferring significant gain (red) or loss (blue) of fitness as determined by the TRANSIT resampling method. Horizontal and vertical dash lines denote the adjusted *p*-value and log_2_-fold change thresholds of 0.05 and ±1, respectively. B) Fitness of each transposon insertion mutant calculated by comparing its expansion factor relative to the rest of the population in the absence and presence of MBX-4132. Select mutants with insertions in genes of interest are noted. Dashed line denotes the line of correlation for untreated and MBX-4132 treated mutants showing comparable fitness values, dotted lines denote cutoff for ±0.2 gain or loss of fitness.

We also found significant losses in the abundance of mutants with insertions in genes associated with oxidative stress response and DNA repair. Notably, mutants in *uvrD1,* which encodes a DNA helicase crucial for both nucleotide excision repair and non-homologous end joining^36^, exhibited a loss of fitness of ∼3 fold (Fig. 5B). Likewise, insertions in *mutT1*, which encodes an 8-oxo-dGTPase and rescues damage to guanine nucleotides caused by oxidative stress^37^, also conferred loss of fitness (Fig. 5B). Previous studies have demonstrated that DNA damage can lead to stalling of RNA polymerase during transcription, and in these cases, the incomplete transcript could result in increased translational pausing and higher demand for *trans*-translation. Indeed, work from the Kreuzer lab has shown that *trans-*translation is important for tolerance to DNA damage in *E. coli*^38–40^. Moreover, oxidized mRNA with 8-oxoguanosine (8-oxoG) lesions has been described to directly contribute to ribosomal stalling^41^. Since mutants in DNA damage repair pathways likely experienced increased reliance on *trans-*translation, their decrease in fitness is consistent with inhibition of *trans*-translaiton by MBX-4132.

In addition, disruptions in a few of the genes responsible for metal homeostasis conferred significant changes in fitness. Strains with mutations in *irtA* and *mymT* displayed significant growth disadvantages when subjected to MBX-4132 (Fig. 5A). Interestingly, mutants with insertions in two of the genes required for siderophore export and recycling, *mmpL4* and *mmpS4*^42,43^, exhibited a weak gain of fitness of just above 2-fold (Fig. 5B). These findings indicate that disruption of metal homeostasis, especially for iron and copper, may affect the anti-mycobacterial activity of MBX-4132.

One limitation of resampling is that transposon insertions near the 5’ and 3’ ends of each gene and those inside intergenic regions are ignored. Therefore, this analysis was largely limited to non-essential genes and would not reveal hypomorphic insertion mutants associated with essential genes, such as *ssr, smpB*, and those encoding for ribosomal proteins, and promoters. Moreover, resampling does not take into account variability among replicates in terms of growth rate or the composition of the input pool. As a result, we sought to address these constraints by calculating expansion factors and assigning fitness values for individual transposon mutants after DMSO and MBX-4132 exposure, using a method described by van Opijnen, et al.^44^. To pinpoint mutants of interest, we subtracted these fitness values of the MBX-4132 treatment condition from those of the DMSO control and considered insertions with differences of > 0.2 as gain of fitness mutants and < -0.2 as loss of fitness mutants (Dataset S2). Using these metrics, we identified mutants in several 50S ribosomal protein genes with changes in fitness (Fig. 5B). Our analysis also confirmed that insertions in *uvrD1* resulted in the loss of fitness in *M. tuberculosis* in the presence of MBX-4132 and revealed that disruption of *uvrC* and *uvrD2* achieved a similar consequence (Fig. 5B). Collectively, our data suggest that alterations in both *trans*-translation activity and metal homeostasis can both modulate MBX-4132 susceptibility.

### Deletion of the *altRP* operon potentiates MBX-4132 activity under high zinc conditions

In *M. tuberculosis*, there are two sets of paralogues of ribosomal protein genes, consisting of four genes encoding for primary ribosomal proteins (PrimRPs, C+) and five genes encoding for alternative ribosomal proteins (AltRPs, C-), differing in that PrimRPs contain cysteine-rich zinc-binding motifs^45,46^. Under zinc-limiting conditions, the cells replace PrimRPs with AltRPs, releasing bound metal ions in favor of other cellular processes^46^. Because transposon insertions in one of the canonical ribosomal protein genes, *rpmG1,* increased fitness of *M. tuberculosis* during MBX-4132 exposure, and those in its AltRibo-encoding paralog, *rpmG2,* decreased fitness (Fig. 5B), we tested whether AltRPs have a role in mediating MBX-4132 susceptibility. We deleted the *altRP* operon (*Rv2055c-Rv2058c*) (Fig. S7A), which expresses four of the five AltRPs, from *M. tuberculosis* H37Rv and tested the susceptibility of the resulting strain (H37Rv Δ*altRP*) in HZMM. The IC_90_ of MBX-4132 against wild-type *M. tuberculosis* was 0.78 μM, whereas that of the Δ*altRP* strain was 0.39 μM (Fig. 6A), indicating that there is a minor contribution of AltRPs to

**Fig 6.**
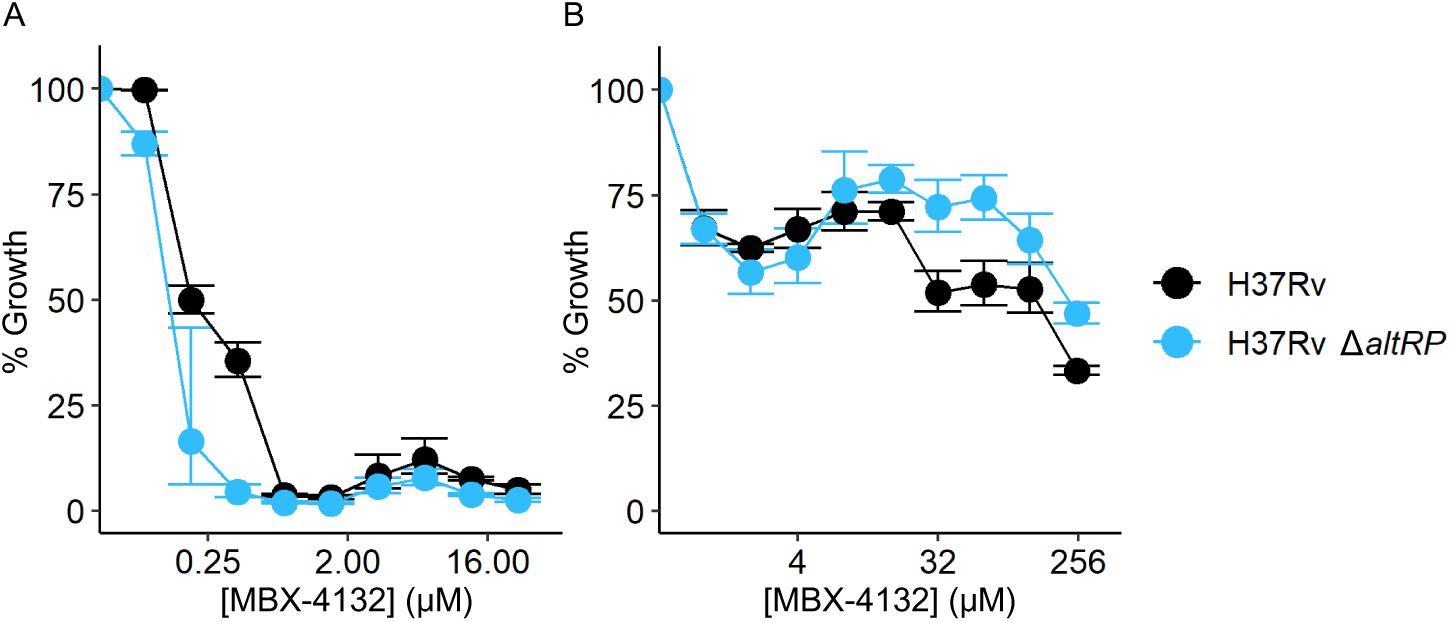
Susceptibility of *M. tuberculosis* H37Rv Δ*altRP* to MBX-4132. *M. tuberculosis* H37Rv (wild type, black) and the Δ*altRP* mutant (light blue) were cultured in (A) HZMM and (B) MM and exposed to MBX-4132 for 14 days. Data represent geometric means and geometric standard deviations for 3 biological replicates.

MBX-4132 susceptibility. We next sought to explore if removing AltRPs confers constitutive susceptibility by testing MBX-4132 against H37Rv Δ*altRP* in MM. We observed that the deletion strain was equally tolerant (IC_50_ = 250 μM, IC_90_ > 250 μM for both strains) (Fig. 6B), indicating that constitutive susceptibility to MBX-4132 cannot be achieved through ribosomes preferentially incorporating PrimRPs over AltRPs. To assess if AltRPs contribute to the activity of MBX-4132 *in vitro*, we repeated the *in vitro trans*-translation reactions with ribosomes purified under zinc-deplete and zinc-rich conditions. However, no significant change in the inhibition of *trans*-translation was observed under either of these conditions (Fig. S7B). Together, these results indicate that the effect of zinc on MBX-4132 activity is not due solely to AltRibos.

### KKL-35 and MBX-4132 are active against *M. tuberculosis* in macrophages

KKL-35 and MBX-4132 were also evaluated in their activities against *M. tuberculosis* H37Rv in RAW 264.7 cells, as a model for *in vivo M. tuberculosis* infections. We observed that 51 μM KKL-35 was able to kill approximately 99% of intracellular *M. tuberculosis* within 3 days of treatment in resting macrophages (Fig. 7A). In contrast, killing by MBX-4132 in resting RAW 264.7 cells was significantly lower, as only around 80% of intracellular bacteria were cleared by 100 μM of the compound (Fig. 7B). Since activated macrophages raise their intracellular zinc concentrations and restrict iron levels as defense mechanisms during bacterial infections^47,48^, we sought to determine whether MBX-4132 would be more potent in interferon (IFN)-γ activated cells. Indeed, IFN-γ activation significantly improved MBX-4132 activity, as shown in the twenty-fold reduction in intracellular bacterial burden by day 7 with 100 μM of MBX-4132 (Fig. 7C).

**Fig 7.**
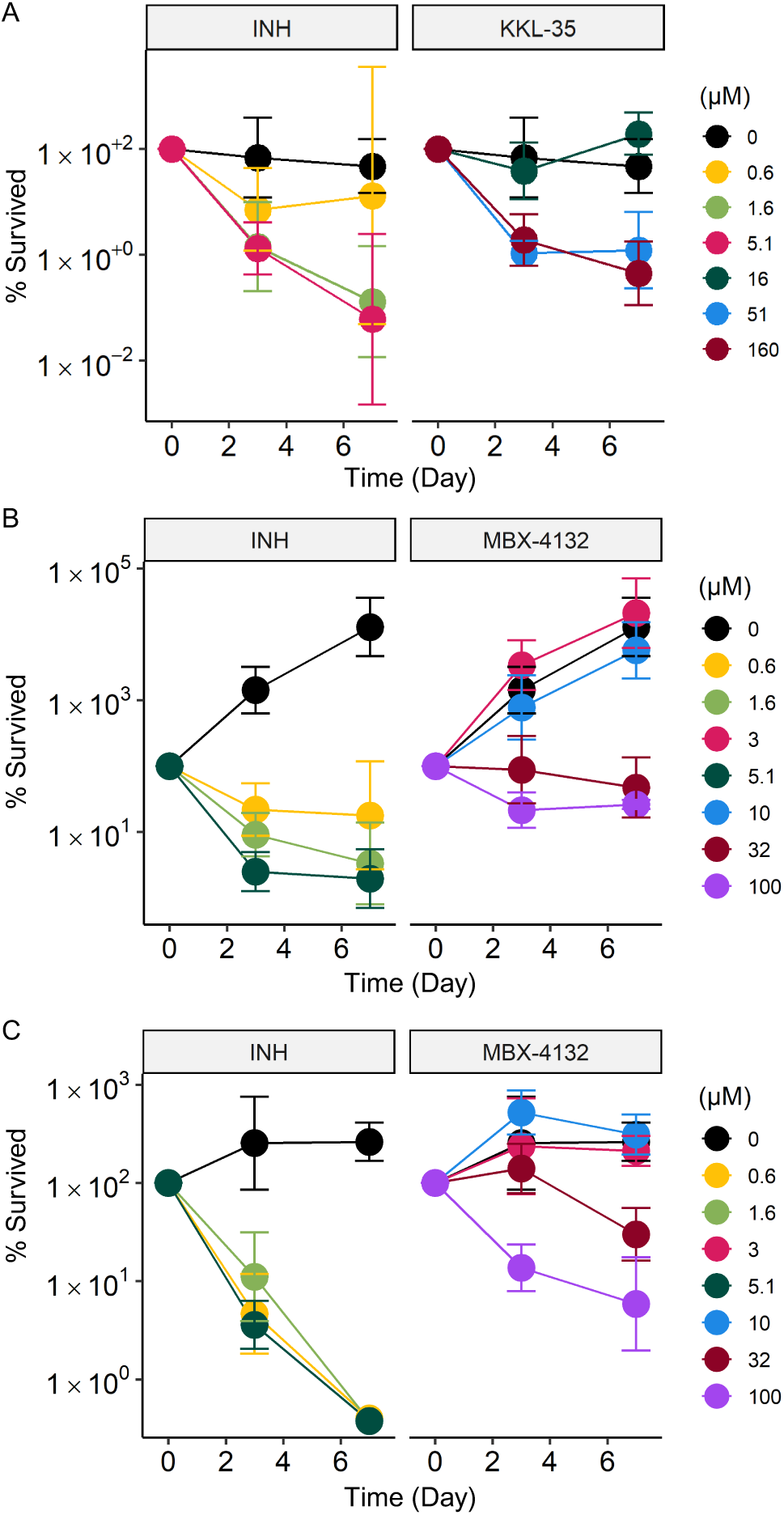
*M. tuberculosis* H37Rv is susceptible to KKL-35 and MBX-4132 killing in RAW 264.7 macrophages. A-B) Resting and C) IFN-γ activated RAW 264.7 macrophages were infected with *M. tuberculosis* H37Rv and subsequently treated with (A) KKL-35 or (B and C) MBX-4132 with INH as a control. Data represent geometric means with error bars indicating the geometric standard deviation for 3 biological replicates.

## DISCUSSION

The activity of MBX-4132 against several mycobacterial species in culture and against *trans-*translation in an *in vitro* system with *M. tuberculosis* components suggests that this acylaminooxadiazole family has promise for anti-mycobacterial therapy. MBX-4132 and KKL-35 are potent and specific inhibitors of *M. tuberculosis trans*-translation *in vitro*. Because *trans*-translation is essential in *M. tuberculosis*, growth inhibition could be due exclusively to inhibition of *trans*-translation. The decreased MIC after depletion of tmRNA is consistent with *trans*-translation being the major target. Similarly, addition of iron ions, which blocks inhibition of *trans*-translation by acylaminooxadiazoles *in vitro*, also blocks growth inhibition by these molecules in culture. In addition, the loss of fitness during exposure to MBX-4132 by mutants with decreasd translation efficiency or fidelity suggests that loss of *trans*-translation activity is a major physiological result of treatment with MBX-4132.

Our data also suggest substantial interactions between acylaminooxadiazoles and metal homeostasis in mycobacteria, including inhibition of acylaminooxadiazole activity by iron and potentiation by zinc. Oxadiazole compounds can complex with transition metals, including iron, zinc, and copper^49,50^. Acylaminooxadiazoles bind the ribosome in a narrow pocket near the peptidyl transfer center^6^, and it is likely that an Fe^3+^- or Fe^2+^-MBX-4132 complex would be too large to bind. Therefore, the effects of iron on MBX-4132 activity may be solely due to preventing MBX-4132 from inhibiting *trans*-translation. On the other hand, potentiation by zinc is difficult to explain through direct interactions with acylaminooxadiazoles, and our data show that the known effect of zinc on ribosomal protein composition has at most a minor contribution to zinc potentiation. Because zinc and iron levels are tightly regulated in *M. tuberculosis*, it is possible that zinc acts by limiting the accessible pool of iron. The ESX-3 secretion system, which is required for siderophore-dependent iron acquisition in *M. tuberculosis*^51,52^, is negatively regulated by high zinc concentration via the zinc-responsive transcription factor Zur^53,54^. In other words, zinc abundance can hinder iron acquisition. Excess zinc also derepresses genes responsible for mycobactin biosynthesis and transport, thus promoting the intracellular accumulation of deferrated siderophores as well as causing an increase in the cells’ demand for iron supply^55,56^. A second possibility is that part of the biological effect of MBX-4132 is through disruption of metal homeostasis. Our transcriptomic data show dysregulation of multiple metal homeostasis pathways after exposure to MBX-4132, including those for iron and copper. Although some of these, mostly iron-independent, responses could be indirect effects of inhibiting *trans*-translation, it is likely that MBX-4132 binds and sequesters intracellular iron, resulting in an iron starvation response. In this case, zinc could exacerbate the effects of iron starvation by decreasing iron uptake and increasing iron utilization through the physiological pathways described above. This model would also explain why disrupting genes involved in iron and copper homeostasis altered the fitness of *M. tuberculosis* after MBX-4132 treatment. Our data also revealed differential expression of genes related to zinc sensing and efflux when tmRNA levels were reduced, suggesting there could be a direct link between *trans*-translation activity and intracellular maintenance of zinc levels. Taken together, these data suggest that disturbing intracellular iron levels could be an additional mechanism of action for MBX-4132.

One previous study examined the interaction between metals and acylaminooxadiazoles with respect to inhibition of *B. subtilis* growth^9^. KKL-40, a close analog of KKL-35, could bind and facilitate transport of copper, and to a lesser extent nickel, across the *B. subtilis* membrane. This interaction leads to synergistic accumulation of copper and KKL-40 inside *B. subtilis* after co-treatment, and a corresponding synergy in growth inhibition. The contrast between inhibition of acylaminooxadiazole activity by iron in mycobacteria and stimulation of acylaminooxadiazole activity by copper in *B. subtilis* may be due in part to the unique properties of the mycobacterial cell wall. The diverse and profound impact of metal ions on acylaminooxadiazole activity highlights the importance of testing potential antibiotics under a variety of physiologically relevant growth conditions.

In this work, KKL-35 showed impressive antitubercular activity in resting RAW 264.7 macrophages. However, pharmacokinetic properties make KKL-35 unsuitable for animal and clinical use. Nonetheless, our KKL-35 data provide an important proof of concept that ribosome rescue is a promising drug target against intracellular *M. tuberculosis* in an infection model. In contrast, MBX-4132 possessed far better pharmacokinetic and bioavailability properties^8^, but its bactericidal activity in macrophages was moderate, even after IFN-γ activation. *M. tuberculosis* encounters diverse microenvironments during infection and can manipulate metal ion trafficking inside host macrophages for survival^57,58^, so the efficacy of MBX-4132 is likely to vary widely in different infection microenvironments. Future efforts should characterize the anti-tubercular efficacy of MBX-4132 in animal models, as well as design compounds that can bypass inhibitory metal interactions.

## METHODS

### Bacterial strains, plasmids, and growth conditions

Bacterial strains, plasmids, and primer sequences are shown in SI Table 1. *M. avium* subspecies avium, *M. abscessus* subspecies abscessus*, M. tuberculosis* H37Rv, and *M. tuberculosis* H37Rv Δ*RD1* Δ*panCD* cells were grown in Middlebrook 7H9 medium (Difco) supplemented with 10% (v/v) OADC enrichment (Difco), 0.2% (v/v) glycerol, 0.05% (v/v) tyloxapol. 50 mg/L pantothenate was added to *M. tuberculosis* Δ*RD1* Δ*panCD* cultures*. Escherichia coli* strain DH5α was used for the propagation of plasmids and strain BL21 (DE3) was used for overexpression and purification of the ten *M. tuberculosis* translation factors and grown in Lysogeny Broth (LB) supplemented with 50 µg/mL kanamycin.

### Overexpression of *M. tuberculosis* translation factors

Plasmids to over-express the 10 *M. tuberculosis* translation factors (IF-1, IF-2,IF-3, EF-G, EF-Tu, EF-Ts, RF-1, RF-2, and RRF) were constructed via HiFi assembly (New England Biolabs). Each gene was amplified from *M. tuberculosis* H37Rv Δ*RD1* Δ*panCD* genomic DNA by PCR and ligated onto the pET28a vector (Addgene) that had been digested with the NcoI and XhoI restriction enzymes (Thermo Fisher). *E. coli* strain DH5α was used for the propagation of plasmids and strain BL21 (DE3) was used for overexpression and purification of protein. *E. coli* BL21 (DE3) strains over-expressing the 10 *M. tuberculosis* translation factors were grown individually in 100 mL Terrific Broth supplemented with 50 µg/mL kanamycin at 37 °C to OD_600_ = 0.6. 6His-tagged *M. tuberculosis* translation factors were overexpressed by growth in the presence of 1 mM isopropyl-thio-β-D-galactoside (IPTG) for 3 h. The ten cultures were pooled, cells were harvested by centrifugation at 6953 *g* for 20 min, resuspended in buffer A (50 mM HEPES-KOH [pH 7.6], 1 M NH_4_Cl, 10 mM MgCl_2,_ 7 mM β-ME), lysed by sonication, and cell debris was removed by centrifugation at 28000 *g* for 20 min. The cleared lysate was incubated with 1 mL HisPur Ni-NTA agarose resin (Thermo Fisher) for 1 h at 4 °C, and washed three times with 40 mL buffer (95% (v/v) buffer A and 5% (v/v) buffer B (50 mM HEPES-KOH [pH 7.6], 100 mM KCl, 10 mM MgCl_2_, 500 mM imidazole, 7 mM β-ME), and bound protein was eluted with 10 mL of elution buffer (10% (v/v) buffer A and 90% (v/v) buffer B). The eluate was dialyzed against buffer (50 mM HEPES-KOH [pH 7.6], 100 mM KCl, 10 mM MgCl_2,_ 30% (v/v) glycerol, and 7 mM β-ME) at 4 °C.

### Overexpression and purification of *M. tuberculosis* EF-Tu

The *tuf* gene was amplified from *M. tuberculosis* H37Rv Δ*RD1* Δ*panCD* genomic DNA by PCR using primers TB_EFTu_F and TB_EFTu_R and ligated into pET28a that had been digested with the NcoI and XhoI. The assembled vector was transformed in *E. coli* BL21-DE3 and this strain was grown in 1 L terrific broth at 37 °C to OD_600_ = 0.6. *M. tuberculosis* EF-Tu was purified and stored as described previously for *E. coli*^59^.

### *M. tuberculosis* protein solution

*M. tuberculosis in vitro* transcription-translation system was designed based on the *E. coli* OnePot PURE system^60^. *M. tuberculosis* protein solution was prepared from the 10 *M. tuberculosis* translation factors as described previously for *E. coli*^60^.

### Energy solution

The energy solution was prepared as described previously^60^. tRNA was extracted and purified from *E. coli* MRE 600 as described previously^61^.

### Ribosome purification from *M. tuberculosis*

H37Rv Δ*RD1* Δ*panCD* cells were grown in Middlebrook 7H9 medium (Difco) supplemented with 10% (v/v) OADC enrichment (Difco), 0.2% (v/v) glycerol, 0.05% (v/v) tyloxapol and pantothenate (50 mg/L) at 37 °C to OD_600_ = 0.35. 1 mM ZnSO_4_ or 1 µM TPEN was supplemented to the medium to induce zinc-rich or zinc-deplete conditions where necessary. Cells were harvested by centrifugation at 6953 *g* for 20 min, resuspended in 30 mL ribosome resuspension buffer (20 mM HEPES-KOH [pH 7.6], 60 mM NH_4_Cl, 12 mM MgCl_2_, 0.5 mM EDTA, 6 mM β-ME), lysed in a French pressure cell, and the lysate was cleared by centrifugation at 28,000 *g* for 30 min at 4 °C. Crude ribosomes were harvested by layering the supernatant over sucrose cushion buffer (37.7% (w/v) sucrose in 20 mM HEPES-KOH [pH 7.6], 500 mM NH_4_Cl, 10 mM MgCl_2_, 0.5 mM EDTA, 6 mM β-ME) followed by centrifugation at 85,000 *g* for 2 h at 4 °C. The ribosome pellet was washed 3 times in ribosome resuspension buffer and resuspended in the same buffer.

### Isolation of *M. tuberculosis* tmRNA and SmpB

*M. tuberculosis* tmRNA was transcribed *in vitro* using Mtb ssrA F and Mtb ssrA R primers (SI methods) based on a previous protocol^62^. *M. tuberculosis* SmpB was overexpressed and purified from *E. coli* BL21(DE3) as described previously^62^.

### *M. tuberculosis in vitro* translation and *trans*-translation assays

Assays were performed as described previously, with some modifications^7^. Translation assays were set up using energy solution (2 µL), *M. tuberculosis* protein solution (1 µL), EF-Tu (10 µM), ribosomes (1.28 µM), DHFR-stop template (9 ng/µL), and [^35^S]-methionine (0.42 µCi/µL). Translation was assayed *in vitro* by expressing full-length DHFR from a DHFR gene with a stop codon (DHFR-stop). DHFR-stop template was prepared via PCR, as described previously^7^. Reactions were incubated at 37 °C for 2.5 h, precipitated with acetone, analyzed by SDS-PAGE, and visualized by phosphor imaging (GE Healthcare, Chicago IL). Relative translation activity in the presence of MBX-4132 or chloramphenicol was calculated with respect to the DMSO-treated control and averaged per reaction from three technical repeats.

*in vitro trans*-translation experiments were set up with the following modifications to the translation assay. *trans*-translation was assayed *in vitro* by expressing full-length DHFR from a DHFR gene without an in-frame stop codon (DHFR-ns). DHFR-ns template was prepared via PCR, as described previously^7^. *M. tuberculosis* tmRNA and SmpB were added to the reactions at final concentrations of 2.75 μM and the reactions were incubated at 37 °C for 2.5 h, precipitated with acetone, analyzed by SDS-PAGE, and visualized by phosphor imaging (GE Healthcare). To inhibit background *trans*-translation activity contributed by tmRNA-SmpB from the ribosomes, 0.5 µM anti-ssrA oligonucleotide was added to the reactions containing no tmRNA-SmpB. To assess the effect of iron and zinc, Fe_2_(SO_4_)_3_ or FeSO_4_ or ZnSO_4_ or water was preincubated with MBX-4132, KKL-35, or DMSO for 10 min at room temperature and added to the *in vitro trans*-translation reactions. Wherever necessary, TPEN or water was preincubated with Fe_2_(SO_4_)_3_ for 10 min at room temperature and added to MBX-4132, KKL-35, or DMSO. This mixture was incubated for 10 min and added to the *in vitro* reactions. Unless stated otherwise the following final concentrations were used: TPEN (150 µM), Fe_2_(SO_4_)_3_ (150 µM), FeSO_4_ (150 µM), ZnSO_4_ (100 µM), MBX-4132 (15 µM), KKL-35 (15 µM). Tagging efficiency was calculated as the percentage of total DHFR tagged by tmRNA-SmpB and averaged per reaction from three experiments. Dose-dependent inhibition of *trans*-translation by MBX-4132 was determined from at least three repeats. GraphPad Prism was used to plot and fit the data to a sigmoidal function and determine the IC_50_.

### MIC and MBC assays

Mtb Minimal media (MM) was prepared as described previously with 0.1% glycerol (v/v)^63^. Low iron minimal media (LIMM) was prepared by omitting ferric ammonium citrate from MM. MIC values were determined using broth dilution assays in 96-well microtiter plates per CLSI guidelines. Plates were incubated at 37 °C (one week for *M. tuberculosis*, four days for *M. avium,* and two days for *M. abscessus*) and the MIC was recorded as the lowest concentration of the compound resulting in no visible growth. 5 μL from wells containing the MIC, 2× MIC, and 4× MIC of each compound was spotted on 7H10 agar plates, and grown at 37 °C. An inhibitor was scored as bactericidal if it resulted in a 99% reduction in the CFU at 2× MIC and no recovered CFU at 4× MIC. MBC was recorded as the lowest concentration resulting in no colony-forming units on the 7H10 agar (Difco) plates. Results were recorded from at least three biological repeats.

### RNA-seq

For the MBX-4132 RNA-seq study, log-phase *M. tuberculosis* H37Rv grown in high zinc minimal media (HZMM, MM supplemented with 3.5 μM ZnSO_4_) was diluted to an OD_600_ of ∼0.1 in the same medium and subjected to 1.2 μM MBX-4132 or equal-volume DMSO in triplicates. Cells were incubated with shaking at 37 °C for another 48 h. For the *ssr* KD RNA-seq study, *M. tuberculosis* H37Rv *ssr* KD and NTC strains were grown to log phase in HZMM containing 50 μg/mL kanamycin. Subsequently, cells were inoculated into HZMM with 100 ng/mL ATc at an OD_600_ of 0.01 in triplicates and induced for 7 days.

To obtain RNA, cells were harvested by centrifugation at 4 °C, and resulting pellets were resuspended in 500 μL TRIzol reagent (Invitrogen) containing 1% polyacryl carrier (Molecular Research Center). Next, cells were lysed by two 1-minute rounds of bead beating (Biospec) at maximum speed. 50 μL of 1-bromo-3-chloropropane was added to the lysed samples, which were subsequently centrifuged to separate phases. Equal-volume ethanol was added to the aqueous phase, after which total RNA was isolated and DNA was removed using the Direct-zol RNA MiniPrep Plus Kit (Zymo Research), and cDNA libraries were generated by SeqCenter using the Stranded Total RNA Prep with Ribo-Zero Plus 563 Microbiome kit (Illumina Inc). Final libraries were sequenced on an Illumina Novaseq platform with 150 bp paired-end reads, generating a total of 12 million reads.

Raw reads were pre-processed using an established pipeline^66^. Differential expression analysis was conducted on the resulting feature count files using the R (v4.3.2) package DESeq2 (v1.34.0)^67^. Enrichment analysis of GO categories was performed using the Database for Annotation, Visualization and Integrated Discovery (DAVID) (v6.9)^68,69^. Analysis outputs were visualized using R packages tidyverse (v2.0.0), ggpubr (v0.6.0), PCAtools (v2.14.0), and EnhancedVolcano (v1.20.0).

To compare gene expression profiles of the two RNA-seq experiments, feature count files were merged, and differential expression was computed for all four conditions using the DESeq2 package with NTC samples as control. PCA was subsequently carried out and visualized using PCAtools. Read counts for transcripts expressed at >1 counts per million in ≥2 replicates were normalized and rescaled between -3 and 3, using the edgeR package (v4.0.16)^70^ in R. A dendrogram was consequently constructed with samples and genes clustering by Pearson correlation, using the pheatmap package (v1.0.12).

### Construction of CRISPRi strains

Individual CRISPRi (Clustered Regularly Interspaced Short Palindromic Repeats interference) mutants were constructed as previously described^71^. Briefly, CRISPRi plasmids were engineered using methods developed by Wong and Rock^72^ using *ssr*-targeting and NTC oligos (Table S1) and plRL58 (Addgene #166886) as the backbone. Successful plasmids confirmed using long-read whole plasmid sequencing at Plasmidsaurus were subsequently co-transformed with plRL19 (Addgene #163634) into electrocompetent *M. tuberculosis* H37Rv generated according to a method previously described^73^. Integration of CRISPRi constructs was confirmed using PCR.

To determine the inhibitory activity of MBX-4132 against the CRISPRi strains, cells were induced for 7 days in MM with 100 ng/mL ATc and then inoculated again at an OD_600_ of 0.01 in 5 mL of MM containing various concentrations of MBX-4132. Resulting cultures were grown for 21 additional days at 37 °C with shaking. OD_600_ of each culture was subsequently measured. Percent growth was calculated as the OD_600_ ratio of the MBX-4132-treated culture to drug-free controls, and IC_90_ was defined as the minimum concentration of MBX-4132 resulting in at least 90% reduction in OD_600_ values.

### Transposon mutagenesis and insertion sequencing

A saturated transposon insertion library in *M. tuberculosis* H37Rv was constructed in MM as part of a previous study^74^. An aliquot of the library was inoculated into 3 liquid cultures in HZMM, each to an OD_600_ of 0.01, which were then incubated at 37 °C with shaking until an OD_600_ of ∼0.32 was reached. At this point, cells were once again diluted to an OD_600_ of 0.01 in HZMM and subjected to either 0.5 μM MBX-4132 or an equal volume DMSO. The remaining, undiluted cells were collected by centrifugation and pellets were frozen at -80 °C to determine input. The MBX-4132- and DMSO-treated cultures were allowed to further grow at 37 °C with shaking to an OD_600_ of ∼0.32, after which they were also spun down and frozen at -80 °C. Once all samples were collected, freezer pellets were thawed at room temperature, and genomic DNA was extracted as previously described^75^. Subsequently, libaries was prepared by the University of Minnesota Genomics Center (UMGC) using procedures also described by Thiede et al.^75^ and sequenced on an AVITI Cloudbreak Low (2x150 PE) platform.

About 250 million total reads were generated. Raw read files were processed using an established pipeline^75,76^. Fitness values for each gene were calculated using two different methods. First, fitness was determined using the resampling method on TRANSIT with default parameters^27^, comparing the relative abundance of mutants in MBX-4132-treated samples against DMSO-treated controls for each gene. Using a separate approach with more emphasis on expansion factors and input library composition, fitness values for individual transposon mutants under each treatment condition were derived as was previously described^44^. Output of both methods was visualized using R packages tidyverse, ggpubr, and ggrepel (v0.9.5).

### Construction of *M. tuberculosis* H37Rv *altRP* deletion strain

Deletion of the *altRP* operon from *M. tuberculosis* H37Rv was achieved using oligonucleotide-mediated recombineering followed by Bxb1 integrase targeting (ORBIT)^77^, using procedures as previously established^75^. Electrocompetent cells expressing pKM461 were electroporated with 1 μg of a targeting oligonucleotide (AGCATGGCCTCGGTAAGTTCCCCGGCTTGCCGGATGCGGGTCATGGGCACAGTGCAGCGCGTCGCTGCCTGGTTTGTACCGTACACCACTGAGACCGCGGTGGTTGACCAGACAAACCGCGGCCGGTGACTTGGCAGTGGGCGGACAAGGGGCACCCTTCCTTCGAAGCTCGGCTTATTGAAAATCAT) and 400 ng of the knockout plasmid pKM464. Recovered cells were plated onto supplemented Middlebrook 7H10 medium containing 50 μg/mL hygromycin B (Corning). The presence of the chromosome-pKM464 junctions was screened for in resulting candidates using Illumina Whole Genome Sequencing (200 Mbp), conducted by SeqCenter. Sequencing reads were aligned to the *M. tuberculosis* H37Rv genome (NC_000962.3) using breseq (v0.38.1)^78^ and visualized using the Integrative Genomics Viewer (IGV) (v2.16.1)^79^.

Following confirmation of *altRP* deletion, susceptibility to MBX-4132 was examined using a modified version of the MTT [3-(4,5-dimethylthiazol-2-yl)-2,5-diphenyltetrazolium bromide] assay^80^. Briefly, mid-log H37Rv and H37Rv Δ*altRP* cultures grown in HZMM or MM were diluted to an OD_600_ = 0.01 in their respective media and then exposed to various concentrations of MBX-4132 at 2.5% volume in a microtiter plate for 14 days. After overnight MTT treatment and formazyn solubilization, the absorbance at 570 nm (OD_570_) of each well was measured using a microplate reader (BioTek Synergy H1). Percent growth, IC_50_, and IC_90_ were calculated as described in the *ssr* KD MBX-4132 susceptibility assay.

### Evaluation of KKL-35 and MBX-4132 in macrophages

*M. tuberculosis* H37Rv was cultured in supplemented Middlebrook 7H9 and HZMM before susceptibility to KKL-35 and MBX-4132 was tested, respectively, to mid-log phase prior to infection.

RAW 264.7 cells were cultured in Dulbecco’s Modified Eagle Medium (DMEM) with GlutaMAX™ supplement (Gibco), 10% fetal bovine serum (FBS), and 100 U/mL of penicillin-streptomycin at 37 °C in a humidified chamber containing 5% CO_2_. Cells were seeded at a density of 10^5^ cells/mL in DMEM with GlutaMAX™ and 10% FBS in 12-well plates. The next day, macrophages were infected as previously outlined and treated with various concentrations of KKL-35, MBX-4132, and isoniazid (INH) (control)^81^. The culture medium was replenished daily. On infection day and days 3 and 7 post-infection, macrophage cultures were lysed with 0.1% Triton X-100 buffer, diluted and plated for CFU/mL quantification. The percentage of cells that survived was calculated as the CFU/mL ratio of the day 3 or 7 culture to its respective day 0 input multiplied by 100%. To examine MBX-4132 killing in activated macrophages, RAW 264.7 cells were treated with fresh DMEM supplemented with GlutaMAX™, 10% FBS, and 5 ng/mL of IFN-γ (Thermo Scientific) the day after seeding and reincubated overnight before infection.

## DATA AVAILABILITY

Sequencing data presented in this study are deposited in the Sequence Read Archive (SRA). BioProject ID: PRJNA1104247

## ACKNOWLEDGEMENTS

We thank William R. Jacobs, Jr. of the Albert Einstein College of Medicine for providing *M. tuberculosis* strains H37Rv and H37Rv Δ*panCD*Δ*RD1*. We are grateful to the NIH NIAID for financial support of this project through grant AI158706. We are also grateful to the University of Minnesota Genomics Center and Biosafety Level 3 Program for providing essential institutional support for this project. We thank Dr. Elise Lamont for her generous advice on macrophage experiements presented in this study.

## SUPPLEMENTAL INFORMATION

**Fig S1.**
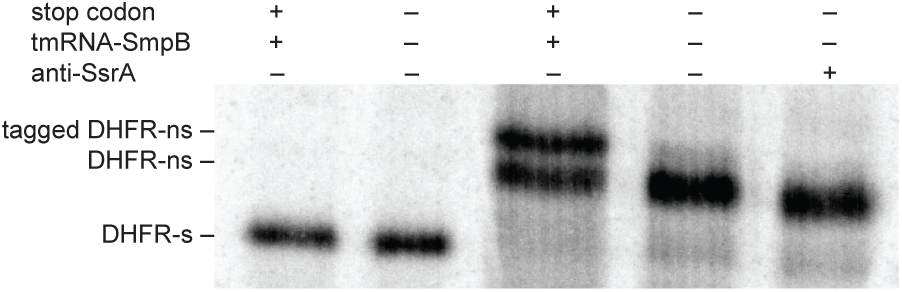
*M. tuberculosis in vitro* translation and *trans*-translation. Genes encoding DHFR with or without a stop codon were expressed *in vitro* in the absence or presence of tmRNA-SmpB and an anti-SsrA oligonucleotide, as indicated. Reactions were incubated at 37 °C for 2.5 h and analyzed by SDS-PAGE followed by phosphorimaging. The locations of DHFR-stop, DHFR-ns, and tagged DHFR as determined in control reactions are indicated.

**Fig S2.**
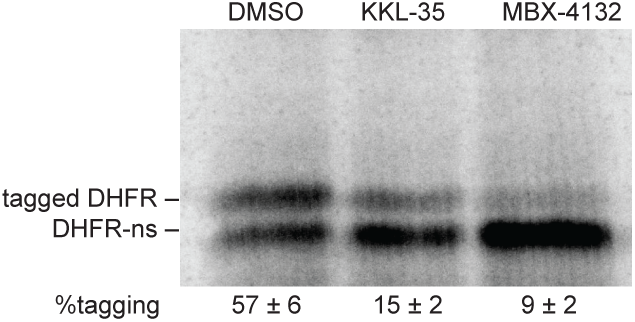
KKL-35 and MBX-4132 inhibit *M. tuberculosis trans*-translation *in vitro.* A gene encoding DHFR without a stop codon was expressed in the presence of *M. tuberculosis* tmRNA-SmpB, with 20 µM KKL-35 or MBX-4132 as indicated. Synthesized protein was detected by incorporation of ^35^S-methionine followed by SDS-PAGE and phosphorimaging. Bands corresponding to tagged and untagged DHFR are indicated, and the average percentage of DHFR protein found in the tagged band for two repeats is shown with the standard deviation.

**Fig S3.**
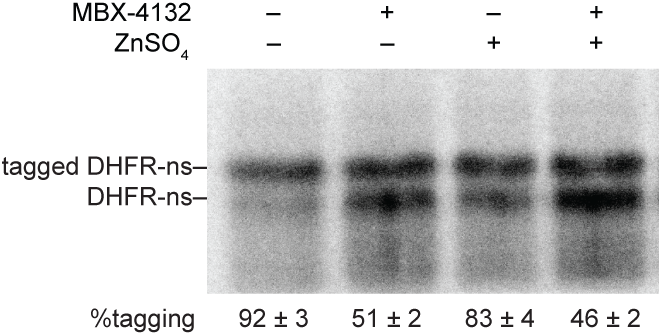
Zinc does not affect the activity of MBX-4132 *in vitro*. A gene encoding DHFR without a stop codon was expressed in the presence of *M. tuberculosis* tmRNA-SmpB without or with 100 µM ZnSO_4_ and 10 µM MBX-4132. Synthesized protein was detected by incorporation of ^35^S-methionine followed by SDS-PAGE and phosphorimaging. Bands corresponding to tagged and untagged DHFR are indicated, and the average percentage of DHFR protein found in the tagged band for two repeats is shown with the standard deviation.

**Fig S4.**
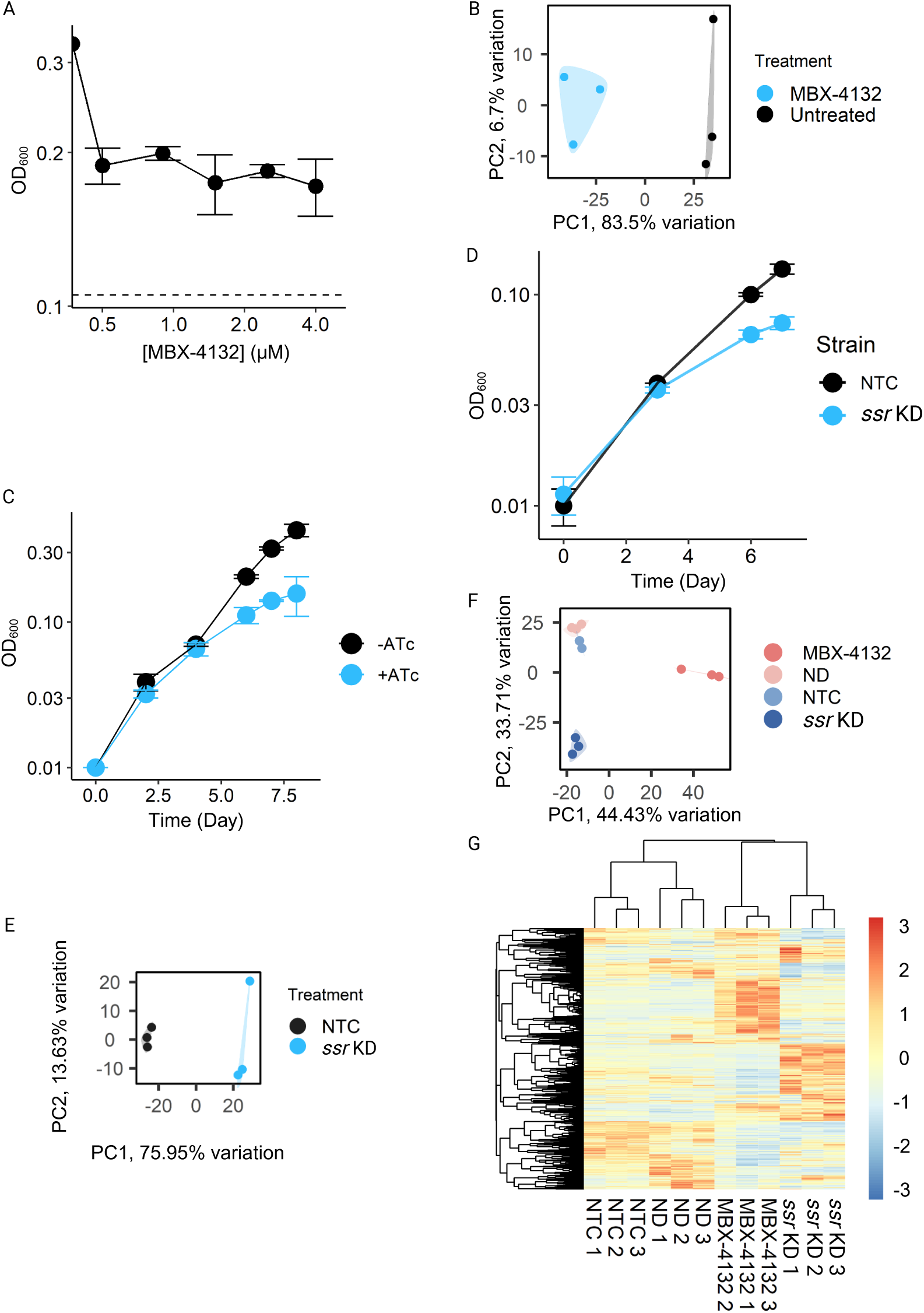
Validation of RNA-seq treatment and results. A) Growth analysis of early-log *M. tuberculosis* H37Rv treated with various concentrations of MBX-4132 for 48 hours (∼1 generation). The dashed line represents OD_600_ of the pre-treatment cultures and the solid line denotes OD_600_ of post-treatment cultures, changing with MBX-4132 concentration. All concentrations tested resulted in ∼50% inhibition of growth, with 1.2 μM selected for RNA-seq as 0.3x IC_90_. B) PCA plot of MBX-4132 RNA-seq results where samples with similar transcriptional profiles cluster. MBX-4132- and DMSO-treated (hereby labeled as Untreated) samples are marked in blue and black, respectively, in 3 biological replicates each. Each point represents a biological replicate. C) Growth validation of the *M. tuberculosis* H37Rv *ssr* KD strain in MM. A mid-log *ssr* KD culture was diluted in MM containing 100 ng/mL ATc to an OD_600_ of ∼ 0.01, and growth of resulting cultures was monitored for 8 days. Data represent means and standard deviations of biological triplicates. D) Growth validation of the *M. tuberculosis* H37Rv *ssr* KD strain in HZMM during RNA-seq setup. E) PCA of the *ssr* KD RNA-seq study. *ssr* KD and NTC samples are shown in light blue and black, respectively, in 3 biological replicates each. F) PCA comparing MBX-4132 and *ssr* KD RNA-seq studies. MBX-4132-treated and DMSO-treated (denoted as No Drug (ND)) were shaded in soft reds, and *ssr* KD and NTC samples were shaded in blues, each containing 3 biological replicates. G) Dendrogram showing expression patterns of genes in MBX-4132 and *ssr* KD RNA-seq studies, with normalized read counts rescaled between -3 and 3. Each row corresponds to a single gene, while each column represents a biological replicate. Genes and samples were clustered using Pearson correlation.

**Figure S5.**
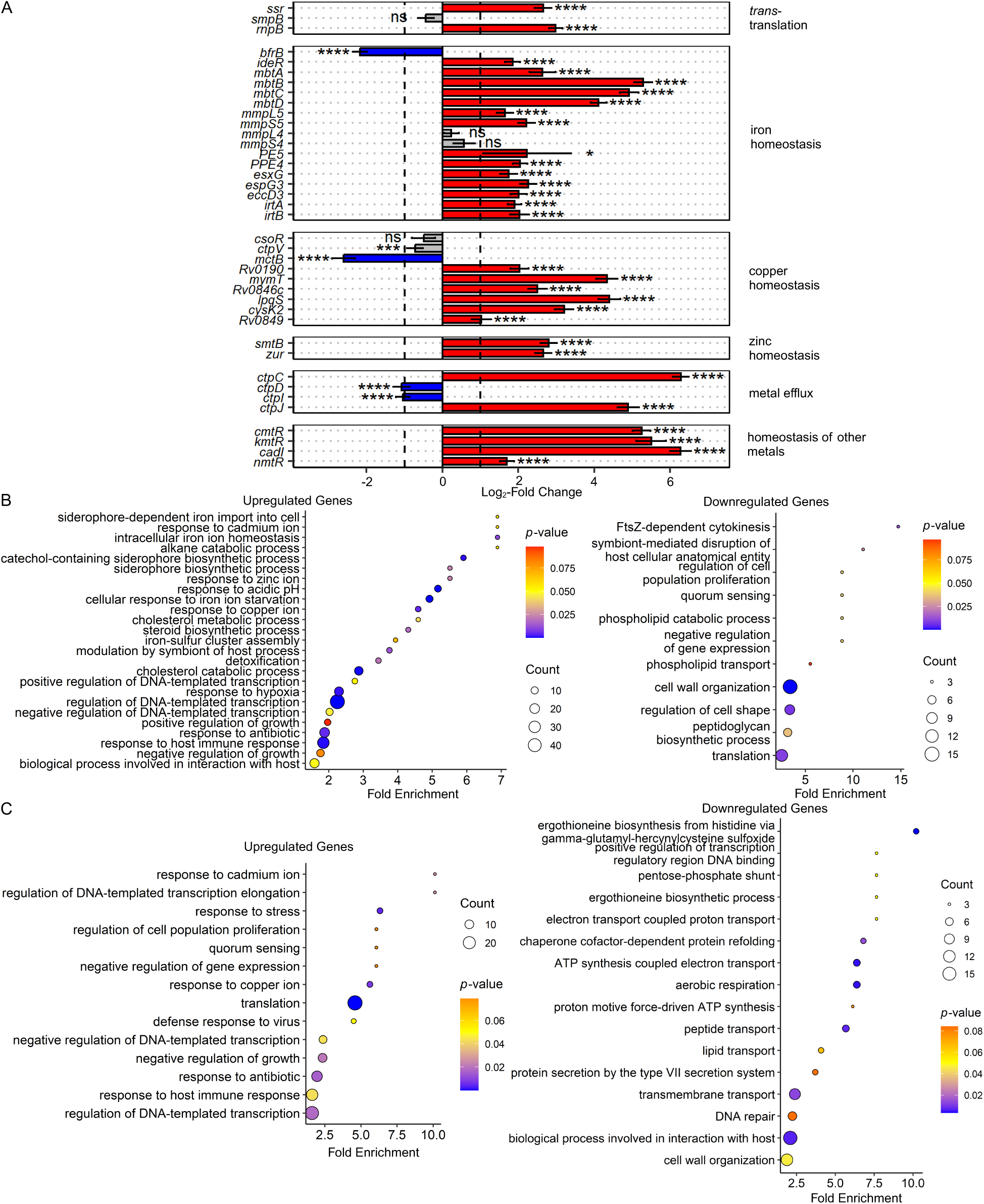
Differential expression of select genes and gene ontology analyses. A) Differential expression of select genes in *M. tuberculosis* H37Rv samples treated with MBX-4132 as compared to DMSO samples. Data represent mean log_2_ fold change ± standard error of the mean of 3 biological replicates. Vertical dashed lines represent log_2_-fold change cut-offs at ±1. Genes are grouped by biological processes. Asterisks indicate statistical significance levels. \**p* < 0.05, \*\**p* < 0.01, \*\*\**p* < 0.001, \*\*\*\**p ≤* 0.001. B-C) GO term enrichment associated with genes significantly up- and down-regulated by (B) MBX-4132 and (C) *ssr* KD, as determined by DAVID.

**Figure S6.**
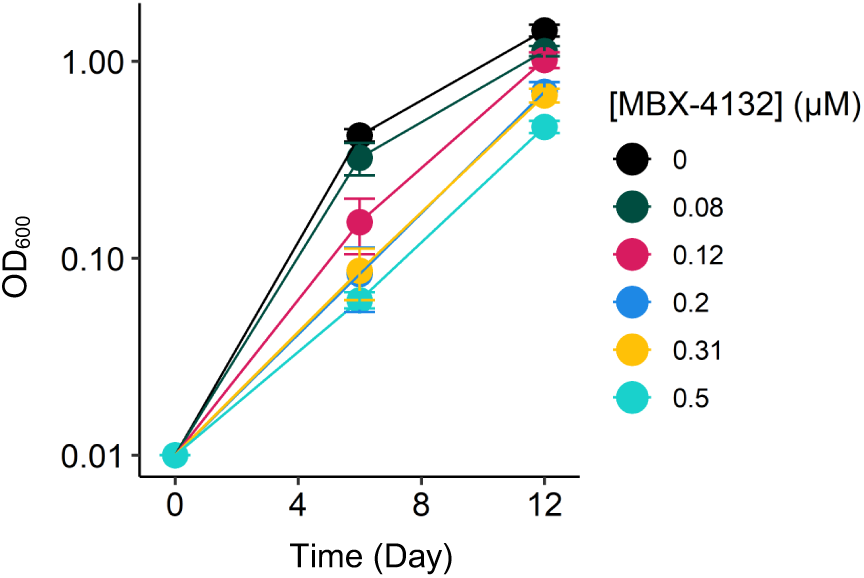
Growth curves of *M. tuberculosis* H37Rv treated with various subinhibitory concentrations of MBX-4132 in HZMM. Mid-log *M. tuberculosis* was diluted to an OD_600_ of 0.01 in HZMM and subjected to a geometric series of MBX-4132 concentrations. OD_600_ of resulting cultures was subsequently measured on days 6 and 12. Different colors represent varying MBX-4132 concentrations, and data depict means of biological triplicates with standard deviations. Treatment of 0.5 μM MBX-4132 approximately doubled the generation time of *M. tuberculosis* H37Rv and was therefore selected as the concentration used for Tn-Seq.

**Figure S7.**
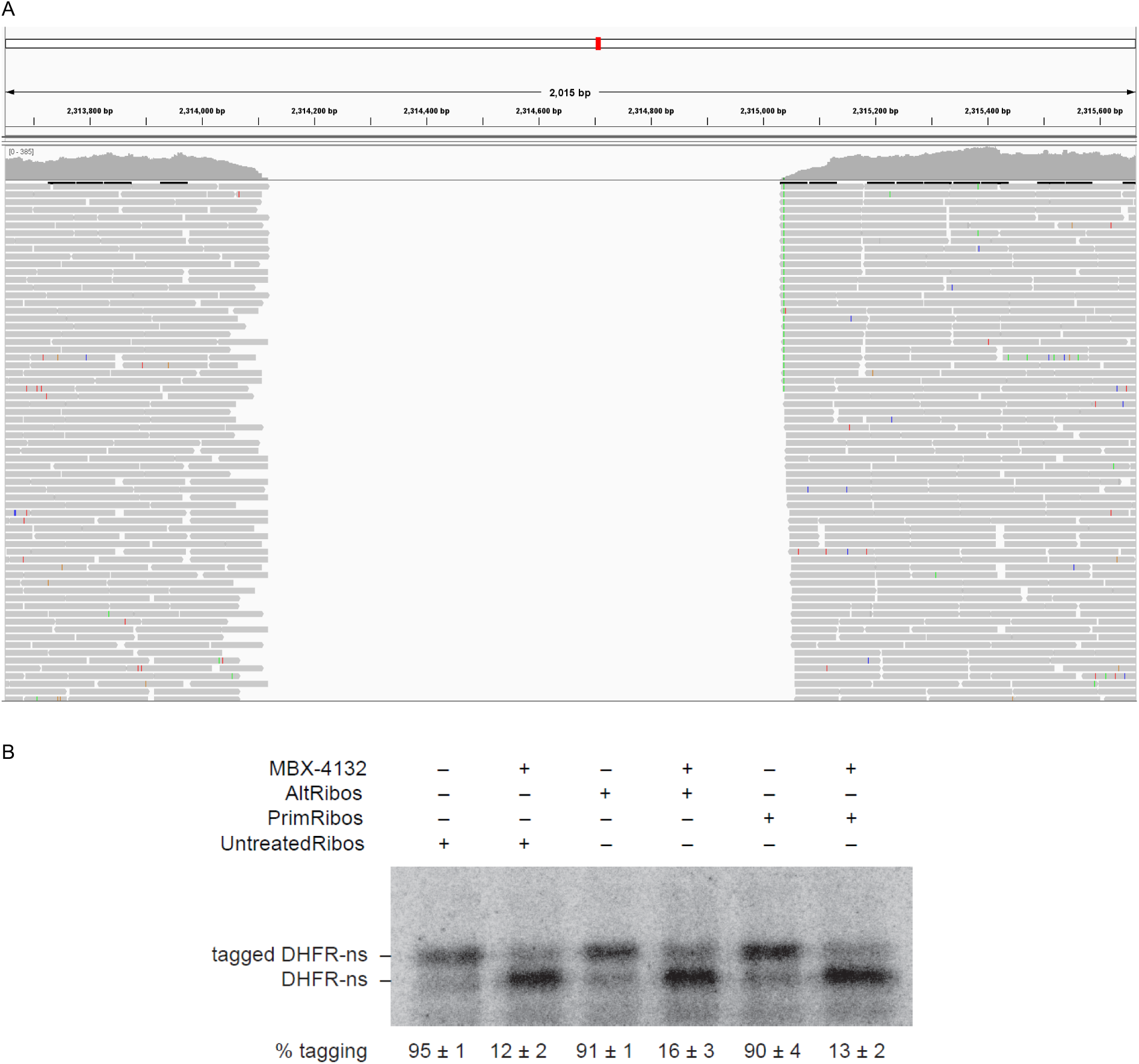
Role of AltRPs in MBX-4132 susceptibility. A) Read alignments of *M. tuberculosis* H37Rv Δ*altRP*. gDNA of the ORBIT deletion strain was extracted, and whole genome resequencing was performed by SeqCenter. Subsequently, reads were aligned to the *M. tuberculosis* H37Rv genome using breseq. BAM files generated were loaded into the Integrative Genomics Viewer (IGV) to illustrate the pileup against the H37Rv genome, zoomed in on a region encapsulating the *altRP* operon (positions 2,313,649-2,315,662). B) AltRPs do not affect acylaminooxadiazole-mediated inhibition of *trans*-translation *in vitro.* A gene encoding DHFR without a stop codon was expressed in the presence of *M. tuberculosis* tmRNA-SmpB and ribosomes purified under zinc-rich (PrimRibos) or zinc deplete conditions (AltRibos). Control reactions were performed in the presence of ribosomes purified from *M. tuberculosis* cultures not supplemented with either ZnSO_4_ or TPEN. 1.3% DMSO or 15 µM MBX-4132 was added to the appropriate reactions. Synthesized protein was detected by incorporation of ^35^S-methionine followed by SDS-PAGE and phosphorimaging. Bands corresponding to tagged and untagged DHFR are indicated, and the average percentage of DHFR protein found in the tagged band for two repeats is shown with the standard deviation.

## SUPPLEMENTAL TABLES

**Table S1.**
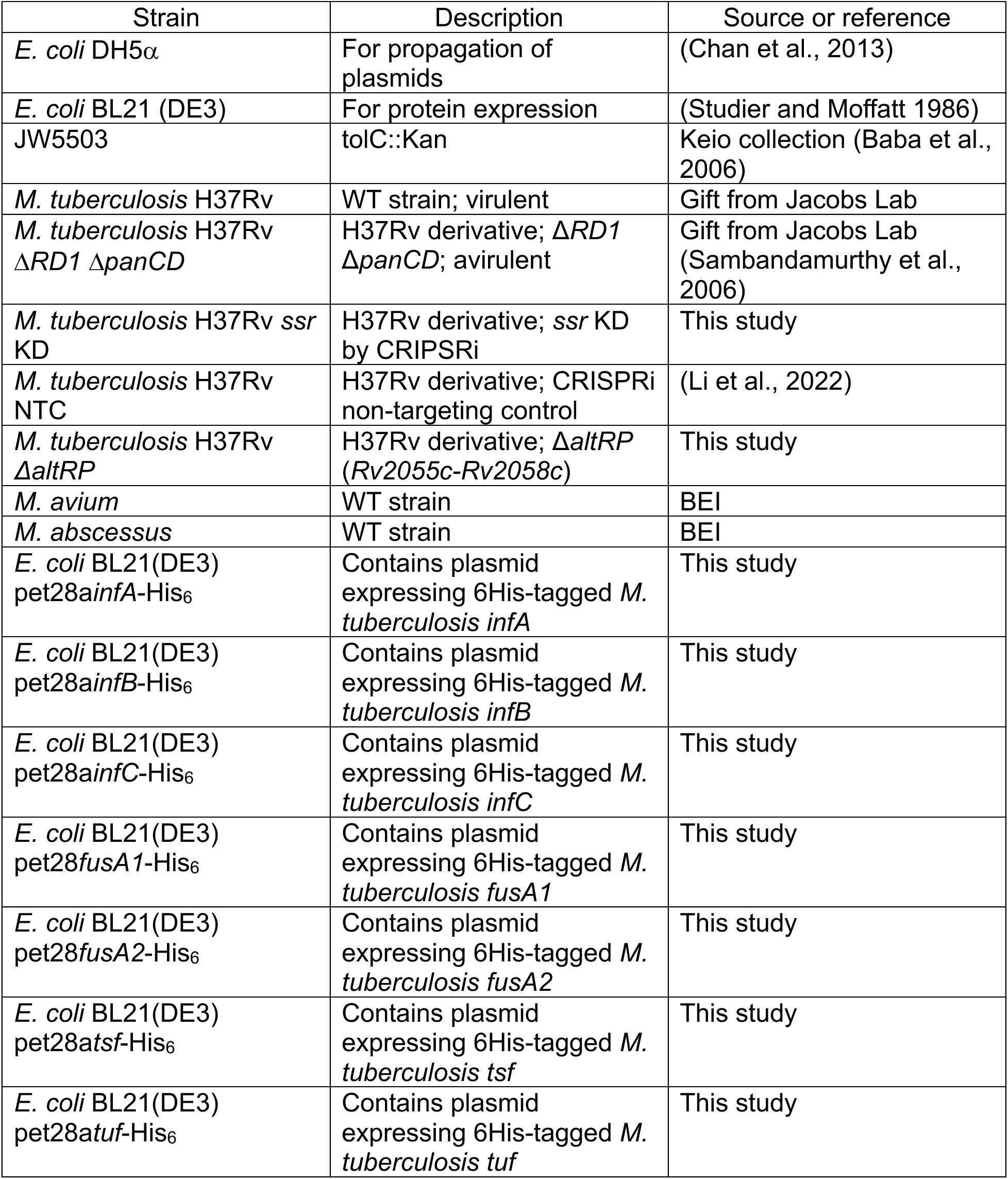
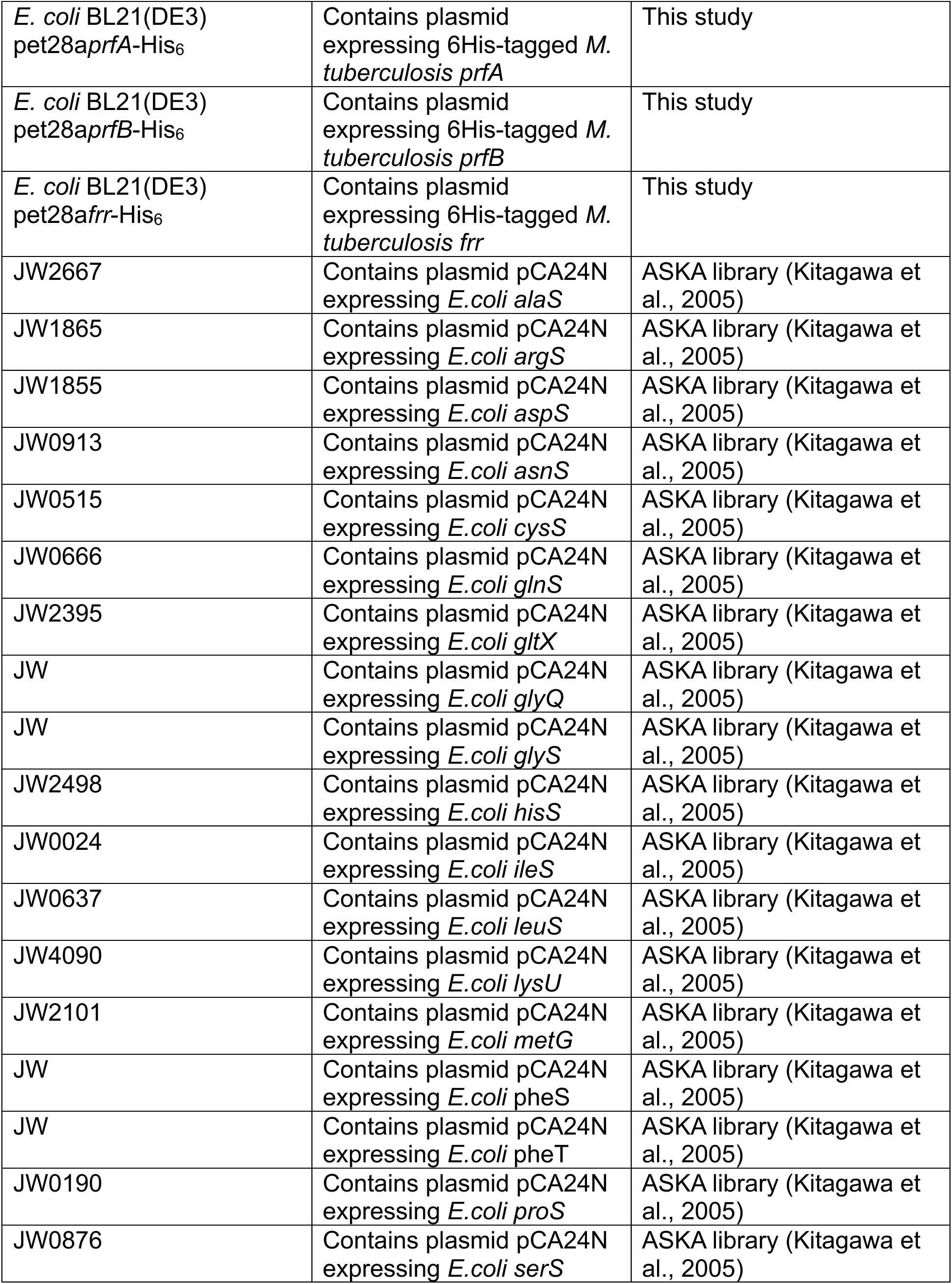
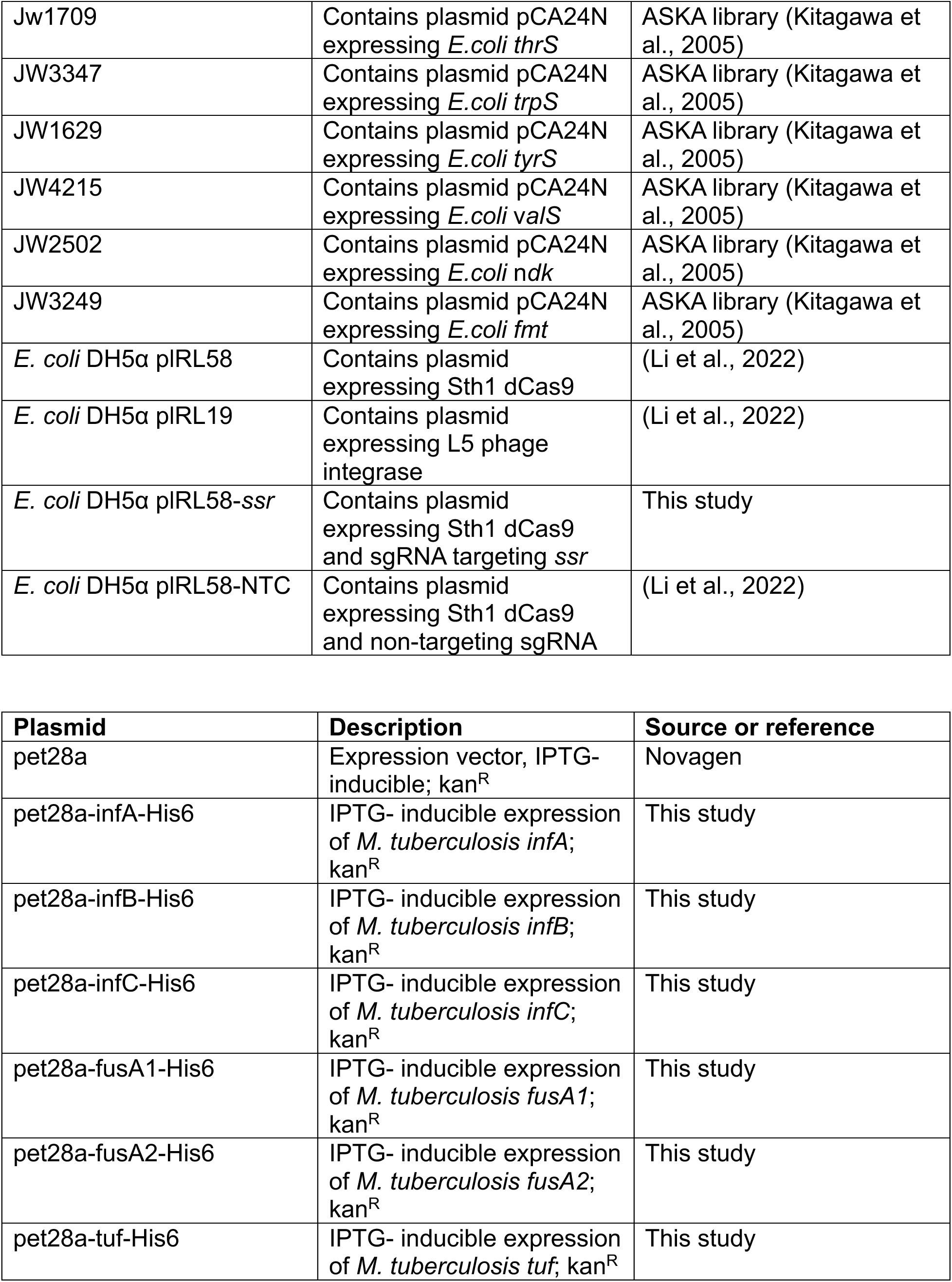
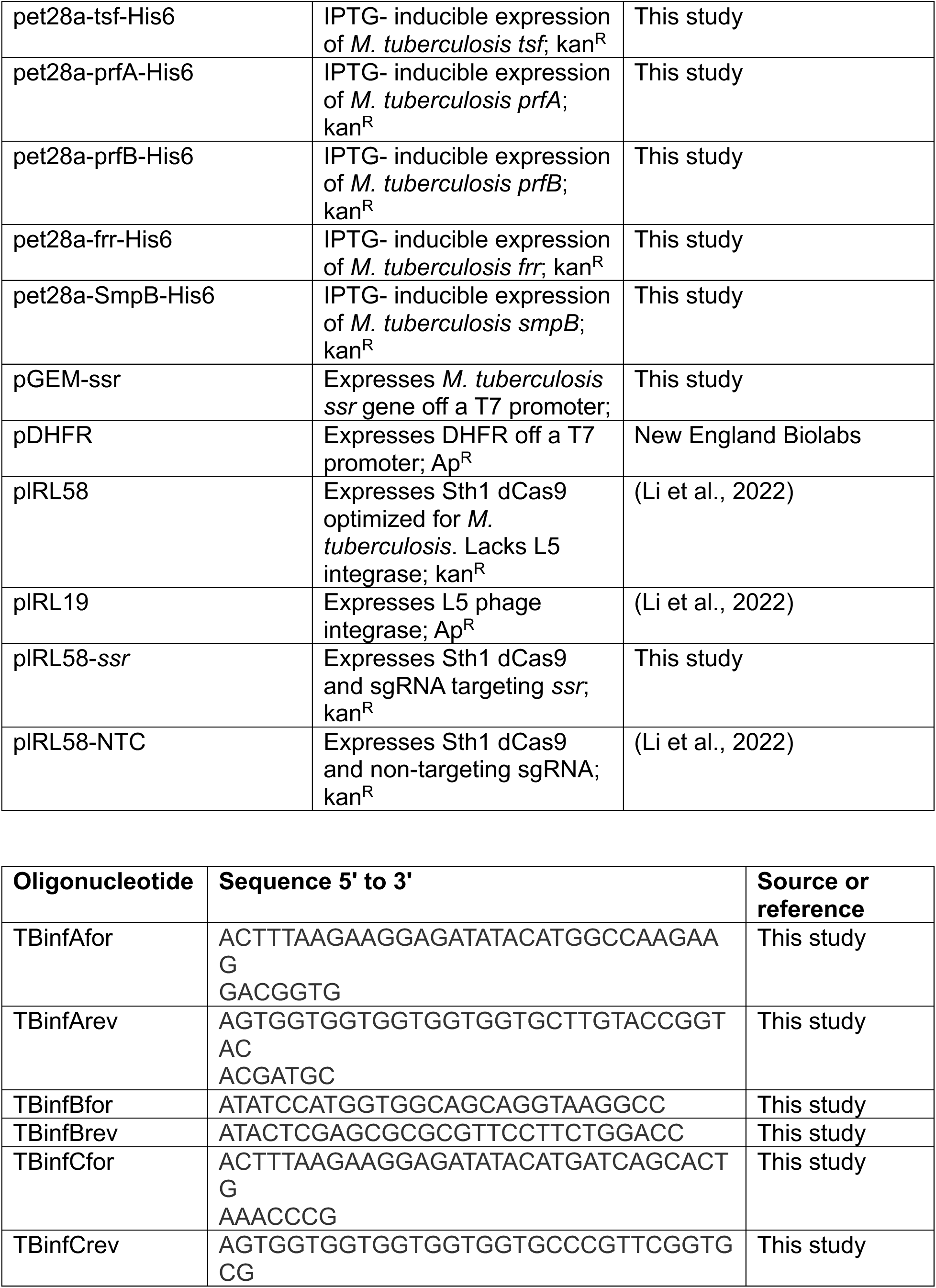
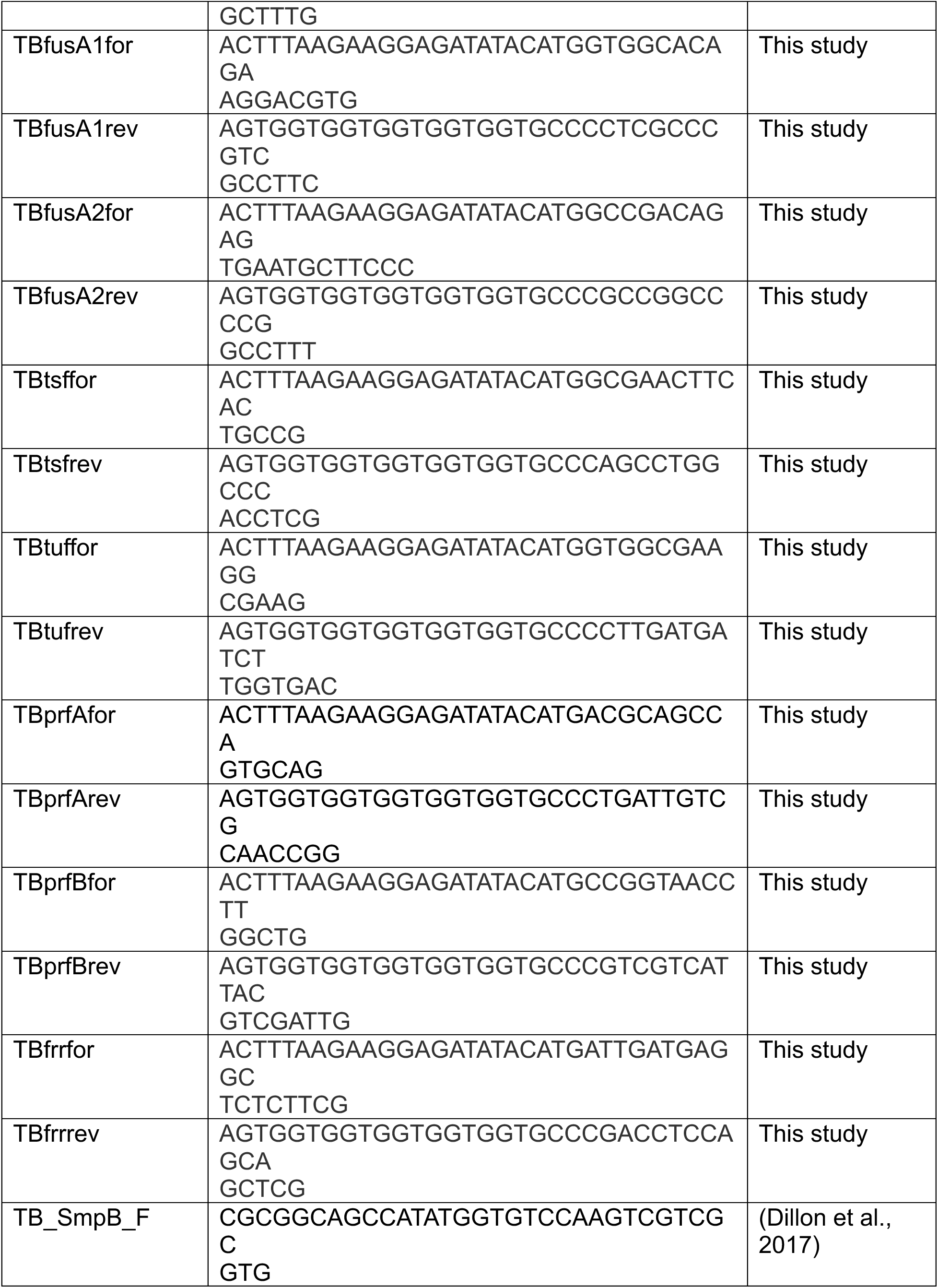
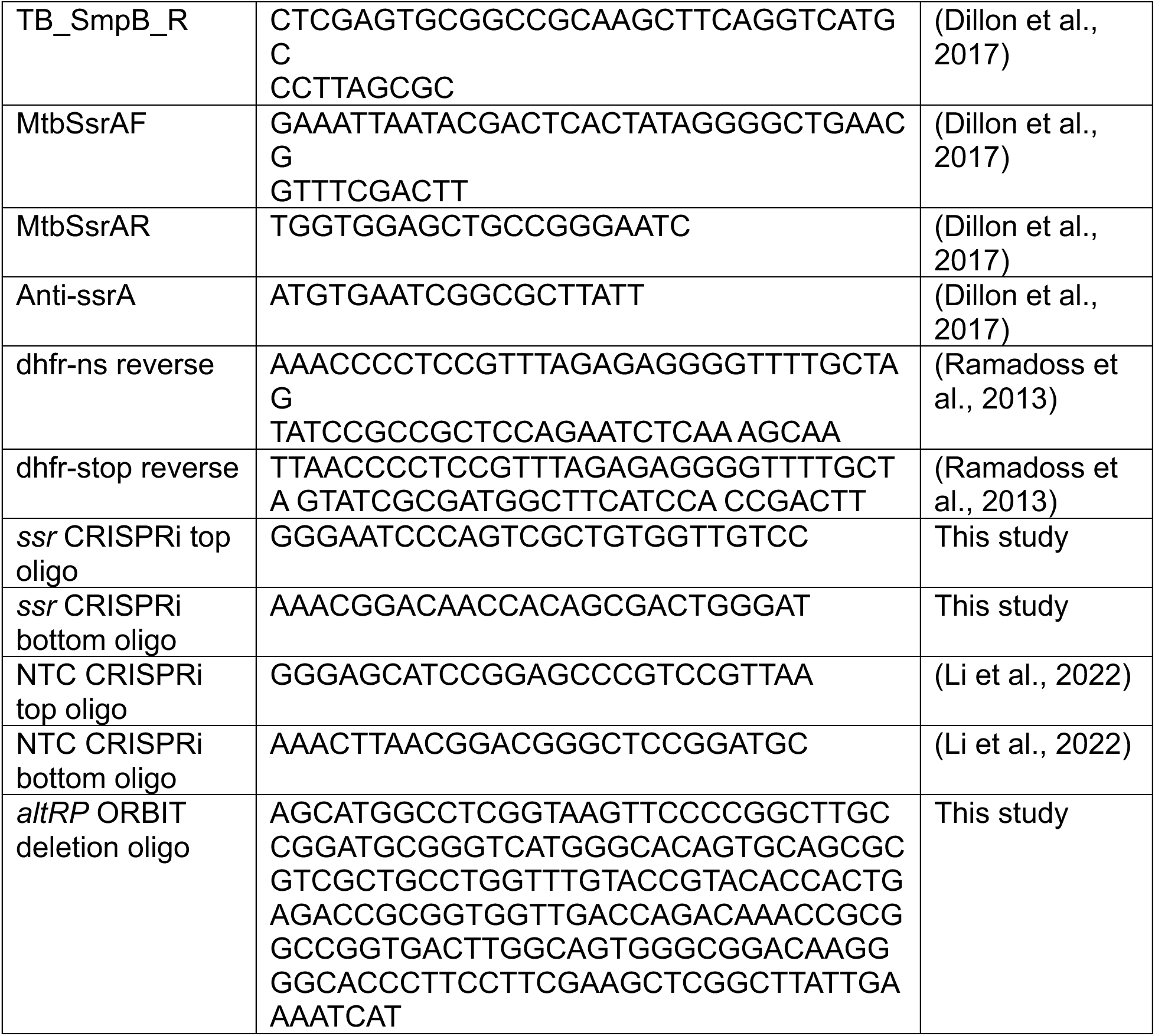
Bacterial strains and plasmids.

## REFERENCES

1. Global tuberculosis report 2023. Geneva: World Health Organization; 2023. Licence: CC BY-NC-SA 3.0 IGO

2. Sekaggya-Wiltshire, C., von Braun, A., Scherrer, A.U., Manabe, Y.C., Buzibye, A., Muller, D., Ledergerber, B., Gutteck, U., Corti, N., Kambugu, A., et al. (2017). Anti-TB drug concentrations and drug-associated toxicities among TB/HIV-coinfected patients. Journal of Antimicrobial Chemotherapy 72, 1172–1177. 10.1093/jac/dkw534.

3. Chakaya, J., Khan, M., Ntoumi, F., Aklillu, E., Fatima, R., Mwaba, P., Kapata, N., Mfinanga, S., Hasnain, S.E., Katoto, P.D.M.C., et al. (2021). Global Tuberculosis Report 2020 – Reflections on the Global TB burden, treatment and prevention efforts. Int J Infect Dis 113, S7–S12. 10.1016/j.ijid.2021.02.107.

4. Keiler, K.C. (2008). Biology of *trans*-Translation. Annu. Rev. Microbiol. 62, 133–151. 10.1146/annurev.micro.62.081307.162948.

5. Keiler, K.C., Waller, P.R.H., and Sauer, R.T. (1996). Role of a Peptide Tagging System in Degradation of Proteins Synthesized from Damaged Messenger RNA. Science 271, 990– 993. 10.1126/science.271.5251.990.

6. Alumasa, J.N., Manzanillo, P.S., Peterson, N.D., Lundrigan, T., Baughn, A.D., Cox, J.S., and Keiler, K.C. (2017). Ribosome Rescue Inhibitors Kill Actively Growing and Nonreplicating Persister Mycobacterium tuberculosis Cells. ACS Infect. Dis. 3, 634–644. 10.1021/acsinfecdis.7b00028.

7. Ramadoss, N.S., Alumasa, J.N., Cheng, L., Wang, Y., Li, S., Chambers, B.S., Chang, H., Chatterjee, A.K., Brinker, A., Engels, I.H., et al. (2013). Small molecule inhibitors of *trans* - translation have broad-spectrum antibiotic activity. Proc. Natl. Acad. Sci. U.S.A. 110, 10282–10287. 10.1073/pnas.1302816110.

8. Aron, Z.D., Mehrani, A., Hoffer, E.D., Connolly, K.L., Srinivas, P., Torhan, M.C., Alumasa, J.N., Cabrera, M., Hosangadi, D., Barbor, J.S., et al. (2021). trans-Translation inhibitors bind to a novel site on the ribosome and clear Neisseria gonorrhoeae in vivo. Nat Commun 12, 1799. 10.1038/s41467-021-22012-7.

9. Senges, C.H.R., Stepanek, J.J., Wenzel, M., Raatschen, N., Ay, Ü., Märtens, Y., Prochnow, P., Vázquez Hernández, M., Yayci, A., Schubert, B., et al. (2020). Comparison of Proteomic Responses as Global Approach to Antibiotic Mechanism of Action Elucidation. Antimicrob Agents Chemother 65, e01373–20. 10.1128/AAC.01373-20.

10. Pandey, A.K., and Sassetti, C.M. (2008). Mycobacterial persistence requires the utilization of host cholesterol. Proceedings of the National Academy of Sciences 105, 4376–4380. 10.1073/pnas.0711159105.

11. Chohan, Z.H., Supuran, C.T., and Scozzafava, A. (2005). Metal binding and antibacterial activity of ciprofloxacin complexes. J Enzyme Inhib Med Chem 20, 303–307. 10.1080/14756360310001624948.

12. Imran, M., Iqbal, J., Iqbal, S., and Ijaz, N. (2007). In Vitro Antibacterial Studies of Ciprofloxacin-imines and Their Complexes with Cu(II),Ni(II),Co(II), and Zn(II). Turkish Journal of Biology 31, 67–72.

13. Patel, M., Chhasatia, M., and Parmar, P. (2010). Antibacterial and DNA interaction studies of zinc(II) complexes with quinolone family member, ciprofloxacin. Eur J Med Chem 45, 439–446. 10.1016/j.ejmech.2009.10.024.

14. Zarkan, A., Macklyne, H.-R., Truman, A.W., Hesketh, A.R., and Hong, H.-J. (2016). The frontline antibiotic vancomycin induces a zinc starvation response in bacteria by binding to Zn(II). Sci Rep 6, 19602. 10.1038/srep19602.

15. Abdeldaim, G., Svensson, E., Blomberg, J., and Herrmann, B. (2016). Duplex detection of the Mycobacterium tuberculosis complex and medically important non-tuberculosis mycobacteria by real-time PCR based on the rnpB gene. APMIS 124, 991–995. 10.1111/apm.12598.

16. Singh, A., Ubaid-ullah, S., Ramteke, A.K., and Batra, J.K. (2016). Influence of Conformation of M. tuberculosis RNase P Protein Subunit on Its Function. PLoS One 11, e0153798. 10.1371/journal.pone.0153798.

17. Lin-Chao, S., Wei, C.-L., and Lin, Y.-T. (1999). RNase E is required for the maturation of ssrA RNA and normal ssrA RNA peptide-tagging activity. Proceedings of the National Academy of Sciences 96, 12406–12411. 10.1073/pnas.96.22.12406.

18. Khare, G., Nangpal, P., and Tyagi, A.K. (2017). Differential Roles of Iron Storage Proteins in Maintaining the Iron Homeostasis in Mycobacterium tuberculosis. PLoS One 12, e0169545. 10.1371/journal.pone.0169545.

19. Rodriguez, G.M., Voskuil, M.I., Gold, B., Schoolnik, G.K., and Smith, I. (2002). ideR, an Essential Gene in Mycobacterium tuberculosis: Role of IdeR in Iron-Dependent Gene Expression, Iron Metabolism, and Oxidative Stress Response. Infect Immun 70, 3371–3381. 10.1128/IAI.70.7.3371-3381.2002.

20. Shi, X., Festa, R.A., Ioerger, T.R., Butler-Wu, S., Sacchettini, J.C., Darwin, K.H., and Samanovic, M.I. (2014). The Copper-Responsive RicR Regulon Contributes to Mycobacterium tuberculosis Virulence. mBio 5, e00876–13. 10.1128/mBio.00876-13.

21. Wang, S., Fang, R., Wang, H., Li, X., Xing, J., Li, Z., and Song, N. (2024). The role of transcriptional regulators in metal ion homeostasis of Mycobacterium tuberculosis. Front. Cell. Infect. Microbiol. 14. 10.3389/fcimb.2024.1360880.

22. Niederweis, M., Wolschendorf, F., Mitra, A., and Neyrolles, O. (2015). Mycobacteria, Metals, and the Macrophage. Immunol Rev 264, 249–263. 10.1111/imr.12265.

23. Rock, J.M., Hopkins, F.F., Chavez, A., Diallo, M., Chase, M.R., Gerrick, E.R., Pritchard, J.R., Church, G.M., Rubin, E.J., Sassetti, C.M., et al. (2017). Programmable transcriptional repression in mycobacteria using an orthogonal CRISPR interference platform. Nat Microbiol 2, 1–9. 10.1038/nmicrobiol.2016.274.

24. Madsen, C.T., Jakobsen, L., Buriánková, K., Doucet-Populaire, F., Pernodet, J.-L., and Douthwaite, S. (2005). Methyltransferase Erm(37) Slips on rRNA to Confer Atypical Resistance in *Mycobacterium tuberculosis*\*. Journal of Biological Chemistry 280, 38942– 38947. 10.1074/jbc.M505727200.

25. Poulton, N.C., DeJesus, M.A., Munsamy-Govender, V., Kanai, M., Roberts, C.G., Azadian, Z.A., Bosch, B., Lin, K.M., Li, S., and Rock, J.M. (2024). Beyond antibiotic resistance: The *whiB7* transcription factor coordinates an adaptive response to alanine starvation in mycobacteria. Cell Chemical Biology 31, 669–682.e7. 10.1016/j.chembiol.2023.12.020.

26. Rudra, P., Hurst-Hess, K.R., Cotten, K.L., Partida-Miranda, A., and Ghosh, P. (2020). Mycobacterial HflX is a ribosome splitting factor that mediates antibiotic resistance. Proceedings of the National Academy of Sciences 117, 629–634. 10.1073/pnas.1906748117.

27. DeJesus, M.A., Ambadipudi, C., Baker, R., Sassetti, C., and Ioerger, T.R. (2015). TRANSIT - A Software Tool for Himar1 TnSeq Analysis. PLOS Computational Biology 11, e1004401. 10.1371/journal.pcbi.1004401.

28. Agris, P.F. (2004). Decoding the genome: a modified view. Nucleic Acids Research 32, 223–238. 10.1093/nar/gkh185.

29. Gibbs, M.R., Moon, K.-M., Chen, M., Balakrishnan, R., Foster, L.J., and Fredrick, K. (2017). Conserved GTPase LepA (Elongation Factor 4) functions in biogenesis of the 30S subunit of the 70S ribosome. Proceedings of the National Academy of Sciences 114, 980– 985. 10.1073/pnas.1613665114.

30. Landwehr, V., Milanov, M., Hong, J., and Koch, H.-G. (2022). The Role of the Universally Conserved ATPase YchF/Ola1 in Translation Regulation during Cellular Stress. Microorganisms 10, 14. 10.3390/microorganisms10010014.

31. Weiss, L.A., and Stallings, C.L. (2013). Essential Roles for Mycobacterium tuberculosis Rel beyond the Production of (p)ppGpp. J Bacteriol 195, 5629–5638. 10.1128/JB.00759-13.

32. Chugh, S., Tiwari, P., Suri, C., Gupta, S.K., Singh, P., Bouzeyen, R., Kidwai, S., Srivastava, M., Rameshwaram, N.R., Kumar, Y., et al. (2024). Polyphosphate kinase-1 regulates bacterial and host metabolic pathways involved in pathogenesis of Mycobacterium tuberculosis. Proceedings of the National Academy of Sciences 121, e2309664121. 10.1073/pnas.2309664121.

33. Sanyal, S., Banerjee, S.K., Banerjee, R., Mukhopadhyay, J., and Kundu, M. (2013). Polyphosphate kinase 1, a central node in the stress response network of Mycobacterium tuberculosis, connects the two-component systems MprAB and SenX3–RegX3 and the extracytoplasmic function sigma factor, sigma E. Microbiology 159, 2074–2086. 10.1099/mic.0.068452-0.

34. Feaga, H.A., Viollier, P.H., and Keiler, K.C. (2014). Release of Nonstop Ribosomes Is Essential. mBio 5, e01916–14. 10.1128/mBio.01916-14.

35. Goralski, T.D.P., Kirimanjeswara, G.S., and Keiler, K.C. (2018). A New Mechanism for Ribosome Rescue Can Recruit RF1 or RF2 to Nonstop Ribosomes. mBio 9, 10.1128/mbio.02436-18. 10.1128/mbio.02436-18.

36. Chadda, A., Jensen, D., Tomko, E.J., Ruiz Manzano, A., Nguyen, B., Lohman, T.M., and Galburt, E.A. (2022). Mycobacterium tuberculosis DNA repair helicase UvrD1 is activated by redox-dependent dimerization via a 2B domain cysteine. Proceedings of the National Academy of Sciences 119, e2114501119. 10.1073/pnas.2114501119.

37. Patil, A.G.G., Sang, P.B., Govindan, A., and Varshney, U. (2013). Mycobacterium tuberculosis MutT1 (Rv2985) and ADPRase (Rv1700) Proteins Constitute a Two-stage Mechanism of 8-Oxo-dGTP and 8-Oxo-GTP Detoxification and Adenosine to Cytidine Mutation Avoidance *. Journal of Biological Chemistry 288, 11252–11262. 10.1074/jbc.M112.442566.

38. Henderson, M.L., and Kreuzer, K.N. (2015). Functions that Protect Escherichia coli from Tightly Bound DNA-Protein Complexes Created by Mutant EcoRII Methyltransferase. PLOS ONE 10, e0128092. 10.1371/journal.pone.0128092.

39. Krasich, R., Wu, S.Y., Kuo, H.K., and Kreuzer, K.N. (2015). Functions that protect *Escherichia coli* from DNA–protein crosslinks. DNA Repair 28, 48–59. 10.1016/j.dnarep.2015.01.016.

40. Kuo, H.K., Krasich, R., Bhagwat, A.S., and Kreuzer, K.N. (2010). Importance of the tmRNA system for cell survival when transcription is blocked by DNA–protein cross-links. Molecular Microbiology 78, 686–700. 10.1111/j.1365-2958.2010.07355.x.

41. Simms, C.L., Hudson, B.H., Mosior, J.W., Rangwala, A.S., and Zaher, H.S. (2014). An Active Role for the Ribosome in Determining the Fate of Oxidized mRNA. Cell Reports 9, 1256–1264. 10.1016/j.celrep.2014.10.042.

42. Jones, C.M., Wells, R.M., Madduri, A.V.R., Renfrow, M.B., Ratledge, C., Moody, D.B., and Niederweis, M. (2014). Self-poisoning of Mycobacterium tuberculosis by interrupting siderophore recycling. Proceedings of the National Academy of Sciences 111, 1945–1950. 10.1073/pnas.1311402111.

43. Wells, R.M., Jones, C.M., Xi, Z., Speer, A., Danilchanka, O., Doornbos, K.S., Sun, P., Wu, F., Tian, C., and Niederweis, M. (2013). Discovery of a Siderophore Export System Essential for Virulence of Mycobacterium tuberculosis. PLOS Pathogens 9, e1003120. 10.1371/journal.ppat.1003120.

44. van Opijnen, T., Bodi, K.L., and Camilli, A. (2009). Tn-seq: high-throughput parallel sequencing for fitness and genetic interaction studies in microorganisms. Nat Methods 6, 767–772. 10.1038/nmeth.1377.

45. Makarova, K.S., Ponomarev, V.A., and Koonin, E.V. (2001). Two C or not two C: recurrent disruption of Zn-ribbons, gene duplication, lineage-specific gene loss, and horizontal gene transfer in evolution of bacterial ribosomal proteins. Genome Biology 2, research0033.1. 10.1186/gb-2001-2-9-research0033.

46. Prisic, S., Hwang, H., Dow, A., Barnaby, O., Pan, T.S., Lonzanida, J.A., Chazin, W.J., Steen, H., and Husson, R.N. (2015). Zinc Regulates a Switch between Primary and Alternative S18 Ribosomal Proteins in Mycobacterium tuberculosis. Mol Microbiol 97, 263–280. 10.1111/mmi.13022.

47. Byrd, T.F., and Horwitz, M.A. (1989). Interferon gamma-activated human monocytes downregulate transferrin receptors and inhibit the intracellular multiplication of Legionella pneumophila by limiting the availability of iron. J Clin Invest 83, 1457–1465.

48. Na-Phatthalung, P., Min, J., and Wang, F. (2021). Macrophage-Mediated Defensive Mechanisms Involving Zinc Homeostasis in Bacterial Infection. Infectious Microbes & Diseases 3, 175. 10.1097/IM9.0000000000000058.

49. Köhler, C., and Rentschler, E. (2016). The First 1,3,4-Oxadiazole Based Dinuclear Iron(II) Complexes Showing Spin Crossover Behavior with Hysteresis. European Journal of Inorganic Chemistry 2016, 1955–1960. 10.1002/ejic.201501278.

50. Salassa, G., and Terenzi, A. (2019). Metal Complexes of Oxadiazole Ligands: An Overview. International Journal of Molecular Sciences 20, 3483. 10.3390/ijms20143483.

51. Siegrist, M.S., Unnikrishnan, M., McConnell, M.J., Borowsky, M., Cheng, T.-Y., Siddiqi, N., Fortune, S.M., Moody, D.B., and Rubin, E.J. (2009). Mycobacterial Esx-3 is required for mycobactin-mediated iron acquisition. Proceedings of the National Academy of Sciences 106, 18792–18797. 10.1073/pnas.0900589106.

52. Zhang, L., Hendrickson, R.C., Meikle, V., Lefkowitz, E.J., Ioerger, T.R., and Niederweis, M. (2020). Comprehensive analysis of iron utilization by Mycobacterium tuberculosis. PLoS Pathog 16, e1008337. 10.1371/journal.ppat.1008337.

53. Serafini, A., Pisu, D., Palù, G., Rodriguez, G.M., and Manganelli, R. (2013). The ESX-3 Secretion System Is Necessary for Iron and Zinc Homeostasis in Mycobacterium tuberculosis. PLOS ONE 8, e78351. 10.1371/journal.pone.0078351.

54. Maciąg, A., Dainese, E., Rodriguez, G.M., Milano, A., Provvedi, R., Pasca, M.R., Smith, I., Palù, G., Riccardi, G., and Manganelli, R. (2007). Global Analysis of the Mycobacterium tuberculosis Zur (FurB) Regulon. Journal of Bacteriology 189, 730–740. 10.1128/jb.01190-06.

55. Li, X., Chen, L., Wang, Y., Guo, X., and He, Z.-G. (2023). Zinc excess impairs Mycobacterium bovis growth through triggering a Zur-IdeR-iron homeostasis signal pathway. Microbiol Spectr 11, e01069–23. 10.1128/spectrum.01069-23.

56. Xu, Z., Wang, P., Wang, H., Yu, Z.H., Au-Yeung, H.Y., Hirayama, T., Sun, H., and Yan, A. (2019). Zinc excess increases cellular demand for iron and decreases tolerance to copper in *Escherichia coli*. Journal of Biological Chemistry 294, 16978–16991. 10.1074/jbc.RA119.010023.

57. Lenaerts, A., Barry III, C.E., and Dartois, V. (2015). Heterogeneity in tuberculosis pathology, microenvironments and therapeutic responses. Immunological Reviews 264, 288–307. 10.1111/imr.12252.

58. Wagner, D., Maser, J., Lai, B., Cai, Z., Barry, C.E., III, Höner zu Bentrup, K., Russell, D.G., and Bermudez, L.E. (2005). Elemental Analysis of Mycobacterium avium-, Mycobacterium tuberculosis-, and Mycobacterium smegmatis-Containing Phagosomes Indicates Pathogen-Induced Microenvironments within the Host Cell’s Endosomal System1. The Journal of Immunology 174, 1491–1500. 10.4049/jimmunol.174.3.1491.

59. Marathe, N., Nguyen, H.A., Alumasa, J.N., Kuzmishin Nagy, A.B., Vazquez, M., Dunham, C.M., and Keiler, K.C. (2023). Antibiotic that inhibits trans-translation blocks binding of EF-Tu to tmRNA but not to tRNA. mBio 14, e0146123. 10.1128/mbio.01461-23.

60. Lavickova, B., and Maerkl, S.J. (2019). A Simple, Robust, and Low-Cost Method To Produce the PURE Cell-Free System. ACS Synth. Biol. 8, 455–462. 10.1021/acssynbio.8b00427.

61. Zubay, G. (1962). The isolation and fractionation of soluble ribonucleic acid. Journal of Molecular Biology 4, 347–356. 10.1016/S0022-2836(62)80015-1.

62. Dillon, N.A., Peterson, N.D., Feaga, H.A., Keiler, K.C., and Baughn, A.D. (2017). Anti-tubercular Activity of Pyrazinamide is Independent of trans-Translation and RpsA. Scientific Reports 7. 10.1038/s41598-017-06415-5.

63. Pandey, A.K., and Sassetti, C.M. (2008). Mycobacterial persistence requires the utilization of host cholesterol. Proc Natl Acad Sci U S A 105, 4376–4380. 10.1073/pnas.0711159105.

64. Jorgensen, J.H., Hindler, J.F., Reller, L.B., and Weinstein, M.P. (2007). New Consensus Guidelines from the Clinical and Laboratory Standards Institute for Antimicrobial Susceptibility Testing of Infrequently Isolated or Fastidious Bacteria. Clinical Infectious Diseases 44, 280–286. 10.1086/510431.

65. Lewis II, J.S. and Clinical and Laboratory Standards Institute (2023). Performance standards for antimicrobial susceptibility testing 33rd ed. (Clinical and Laboratory Standards Institute).

66. MDHowe4 (2022). MDHowe4/RNAseq-Pipeline.

67. Love, M.I., Huber, W., and Anders, S. (2014). Moderated estimation of fold change and dispersion for RNA-seq data with DESeq2. Genome Biology 15, 550. 10.1186/s13059-014-0550-8.

68. Huang, D.W., Sherman, B.T., and Lempicki, R.A. (2009). Bioinformatics enrichment tools: paths toward the comprehensive functional analysis of large gene lists. Nucleic Acids Research 37, 1–13. 10.1093/nar/gkn923.

69. Huang, D.W., Sherman, B.T., and Lempicki, R.A. (2009). Systematic and integrative analysis of large gene lists using DAVID bioinformatics resources. Nat Protoc 4, 44–57. 10.1038/nprot.2008.211.

70. Robinson, M.D., McCarthy, D.J., and Smyth, G.K. (2010). edgeR: a Bioconductor package for differential expression analysis of digital gene expression data. Bioinformatics 26, 139– 140. 10.1093/bioinformatics/btp616.

71. Li, S., Poulton, N.C., Chang, J.S., Azadian, Z.A., DeJesus, M.A., Ruecker, N., Zimmerman, M.D., Eckartt, K.A., Bosch, B., Engelhart, C.A., et al. (2022). CRISPRi chemical genetics and comparative genomics identify genes mediating drug potency in Mycobacterium tuberculosis. Nat Microbiol 7, 766–779. 10.1038/s41564-022-01130-y.

72. Wong, A.I., and Rock, J.M. (2021). CRISPR Interference (CRISPRi) for Targeted Gene SilencingGene silencing in Mycobacteria. In Mycobacteria Protocols, T. Parish and A. Kumar, eds. (Springer US), pp. 343–364. 10.1007/978-1-0716-1460-0_16.

73. Murphy, K.C., Papavinasasundaram, K., and Sassetti, C.M. (2015). Mycobacterial Recombineering. In Mycobacteria Protocols, T. Parish and D. M. Roberts, eds. (Springer), pp. 177–199. 10.1007/978-1-4939-2450-9_10.

74. Minato, Y., Gohl, D.M., Thiede, J.M., Chacón, J.M., Harcombe, W.R., Maruyama, F., and Baughn, A.D. (2019). Genomewide Assessment of Mycobacterium tuberculosis Conditionally Essential Metabolic Pathways. mSystems 4, e00070–19. 10.1128/mSystems.00070-19.

75. Thiede, J.M., Dillon, N.A., Howe, M.D., Aflakpui, R., Modlin, S.J., Hoffner, S.E., Valafar, F., Minato, Y., and Baughn, A.D. (2022). Pyrazinamide Susceptibility Is Driven by Activation of the SigE-Dependent Cell Envelope Stress Response in Mycobacterium tuberculosis. mBio 13, e00439–21. 10.1128/mbio.00439-21.

76. MDHowe4 (2022). MDHowe4/Himar1-TnSeq-Pipeline.

77. Murphy, K.C., Nelson, S.J., Nambi, S., Papavinasasundaram, K., Baer, C.E., and Sassetti, C.M. (2018). ORBIT: a New Paradigm for Genetic Engineering of Mycobacterial Chromosomes. mBio 9, 10.1128/mbio.01467-18. 10.1128/mbio.01467-18.

78. Deatherage, D.E., and Barrick, J.E. (2014). Identification of Mutations in Laboratory-Evolved Microbes from Next-Generation Sequencing Data Using breseq. In Engineering and Analyzing Multicellular Systems: Methods and Protocols, L. Sun and W. Shou, eds. (Springer), pp. 165–188. 10.1007/978-1-4939-0554-6_12.

79. Robinson, J.T., Thorvaldsdóttir, H., Winckler, W., Guttman, M., Lander, E.S., Getz, G., and Mesirov, J.P. (2011). Integrative genomics viewer. Nat Biotechnol 29, 24–26. 10.1038/nbt.1754.

80. Martin, A., Morcillo, N., Lemus, D., Montoro, E., da Silva Telles, M.A., Simboli, N., Pontino, M., Porras, T., León, C., Velasco, M., et al. (2005). Multicenter study of MTT and resazurin assays for testing susceptibility to first-line anti-tuberculosis drugs. The International Journal of Tuberculosis and Lung Disease 9, 901–906.

81. Cole, M.S., Howe, M.D., Buonomo, J.A., Sharma, S., Lamont, E.A., Brody, S.I., Mishra, N.K., Minato, Y., Thiede, J.M., Baughn, A.D., et al. (2022). Cephem-Pyrazinoic Acid Conjugates: Circumventing Resistance in Mycobacterium tuberculosis. Chemistry – A European Journal 28, e202200995. 10.1002/chem.202200995.

