## Supplementary material for "A *trans*-translation inhibitor is potentiated by zinc and kills *Mycobacterium tuberculosis* and non-tuberculous mycobacteria": Note S1

Transcriptomic responses of *M. tuberculosis* to MBX-4132 were examined in an RNA sequencing (RNA-seq) study (Dataset S1). 1.2 µM MBX-4132 was chosen because that concentration inhibited 50% of bacterial growth over the 48 h treatment window, which is roughly one generation time in high zinc Mtb Minimal Medium (HZMM) (Fig. S4A). This growth medium was chosen over Mtb Minimal Medium (MM) as MBX-4132 was inactive in the latter, whereas low iron MM was not selected due to difficulties in obtaining sufficient cells in balanced growth.

tmRNA levels increased >4-fold in cultures treated with MBX-4132, consistent with cells sensing a deficit in *trans*-translation activity and up-regulating the amount of tmRNA (Fig. 4B & S5A). In contrast, the *smpB* transcript did not change significantly (Fig. 4B & S5A). The observed difference between tmRNA and *smpB* may be due in part to the fact that tmRNA is an RNA, and therefore RNA-seq measured the final gene product and not new transcription. It is possible that *smpB* transcription was up-regulated early after MBX-4132 treatment, and the SmpB protein levels were increased at 48 h even though the transcript was no longer more abundant. Likewise, the RNA component of RNase P, encoded by *rnpB*[1], was more abundant after MBX-4132 treatment, but *rnpA*, the transcript encoding the protein subunit of RNase P[2], did not appear to be up-regulated (Fig. 4B & Dataset S1). RNase P participates in the maturation of tRNA and tmRNA via the removal of the 5'-leader[3], so increased tmRNA and RNase P RNA levels may both result from cells attempting to increase the amount of *trans*-translation in response to inhibition.

Genes involved in metal homeostasis were also differentially regulated after MBX-4132 treatment. Most prominently, transcripts of genes involved in iron uptake and storage were significantly altered. *bfrB*, which encodes ferritin and is crucial for bacterial defense against excess iron, was strongly repressed[4] (Fig. 4B & S5A). Furthermore, nearly all genes involved in siderophore biosynthesis (*mbt* gene cluster, *mbtA-mbtN*)[5] were induced, except *mbtN* (Fig. 4B & S5A). The expression of genes responsible for export of deferrated siderophores (*mmpL5* and *mmpS5*)[6,7], import of ferric-bound siderophores through the outer membrane (ESX-3 operon, including *PE5, PPE4, esxG, espG3, eccD3*)[7], and shuttling of siderophores through the inner membrane back to the cytoplasm and the subsequent reduction and release of bound iron (*irtAB)*[8] was increased (Fig. 4B & S5A). The expression of *ideR*, the iron-sensing transcriptional regulator in *M. tuberculosis*, was also significantly increased (Fig. 4C). In the presence of replete iron, IdeR binds to regions in the genome upstream of numerous genes involved in iron homeostasis, activating or repressing their expression to prevent iron toxicity[9]. Of these, genes including the *mbt* cluster, the ESX-3 operon, and *irtAB* are repressed by high iron and IdeR, whereas *bfrB* is induced[10]. Thus, the responses of these genes after MBX-4132 treatment are similar to those when *M. tuberculosis* are sensing insufficient intracellular iron.

Other metal homeostasis genes were differentially regulated in *M. tuberculosis* treated with MBX-4132, but in a less coherent manner than for iron. Three copper-response pathways have been identified in *M. tuberculosis*, the first being the RicR (*Rv0190*) regulon[11]. A majority of annotated member genes, including *ricR* itself, *mmcO* (*Rv0846c)*, *mymT, lpqS, cysK2,* and *Rv0849* were induced after the addition of MBX-4132 (Fig. 4B & S5A). RicR represses its regulon in the absence of copper, so induction of these genes is consistent with bacteria sensing excess copper[11,12]. In contrast, the four-gene *csoR* operon (*csoR, ctpV*, *Rv0968,* and *Rv0970*)[11] did not change more than 2-fold (Fig. S5A & Dataset S1). Of these genes, *ctpV* expresses an efflux pump which exports excess intracellular copper[13]. Similarly, *mctB*, which is not controlled by RicR or CsoR and encodes an outer membrane protein responsible for removing excess cuprous ions from the cell[14], was significantly repressed after treatment with MBX-4132 (Fig. 4B & S5A). These results, particularly the lack of upregulation of *ctpV* and *mctB*, suggest specific inactivation of RicR instead of an increase of copper in the cell.

Two zinc-sensing transcriptional regulators, *smtB* (*Rv2358*) and *zur*[15], were strongly induced (Fig. 4B & S5A). In the presence of high intracellular zinc concentrations, the repression of *zur* by Rv2358 is reduced, and Zur binds to the operators of genes in the Zur regulon, resulting in repression in its expression[16]. In our study, no transcriptional changes meeting the 2-fold threshold were observed in any genes in the Zur regulon, except those also regulated by IdeR (Fig. 4B & S5A & Dataset S1). Similarly, genes expressing metal-sensing proteins such as *kmtR, cmtR, cadI,* and *nmtR* were strongly induced (Fig. 4B & S5A). These proteins are capable sensing transition metals such as nickel, cobalt, and cadmium[17], but the cognate regulons of these repressors were not repressed. Moreover, several, although not all, genes encoding for metal efflux pumps were differentially expressed[18]. While some were drastically upregulated, such as *ctpC*[19] and *ctpJ*[20], others were repressed, such as *ctpD*[20] and *ctpI*.

Overall, these observations suggest that both *trans*-translation and metal homeostasis were perturbed when *M. tuberculosis* was treated with MBX-4132. However, it is unknown whether the apparent impacts on metals were directly connected to inhibition of *trans*-translation or off-target side effects. Therefore, we constructed a CRISPRi (Clustered Regularly Interspaced Short Palindromic Repeats interference)[21,22] hypomorphic knockdown (KD) strain of *ssr* and a non-targeting control (NTC) strain. The KD strain exhibited slower growth in both HZMM and MM and susceptibility to MBX-4132 in MM after 6-7 days of induction, as compared to the NTC controls (Fig. 4C&D & S4C). This synergy between MBX-4132 treatment and genetic impairment of *trans*-translation suggests that this cellular process is the principal target of MBX-4132 against *M. tuberculosis in vivo*.

RNA-seq was conducted comparing the *ssr* KD strain with the NTC strain after 7 days of induction (Dataset S1). Most notably, tmRNA levels were reduced by 165-fold, which provided further validation for the KD system (Fig. 4E). Repressing tmRNA production and MBX-4132 exposure both led to the induction of genes that are also induced by transcriptional and translational pausing, such as *whiB6*, *whiB7, eis, erm(37),* and *hflX*[23–25] (Fig. 4E & Dataset S1). Numerous genes encoding for amino acid biosynthesis and ribosomal proteins were also significantly upregulated, indicating that *M. tuberculosis* sensed an increased need for active translation due to not having enough ribosomes participating in this crucial cellular process (Fig. 4E & Dataset S1).

Next, the two RNA-seq studies were directly compared to each other. Principle component analysis (PCA) of both studies together revealed that NTC and dimethyl sulfoxide (DMSO)-treated samples clustered closely with one another (Fig. S4F), suggesting that the presence of the CRISPRi system, without targeting any gene, or treatment of a small volume of DMSO do not lead to significant changes in gene expression profiles in *M. tuberculosis*. MBX-4132-treated and *ssr* KD samples appeared different from one another, as inferred by PCA (Fig. S4F), which might be untrue considering the arbitrariness of unit vectors[26]. Clustered heatmaps, on the other hand, displayed that MBX-4132 and *ssr* KD samples clustered more closely to each other than to their respective controls, and numerous genes exhibited similar differential expression in both studies (Fig. S4G). These results are supportive of MBX-4132 possessing on-target activity as a *trans*-translation inhibitor.

We found 109 coding sequences induced and 47 repressed in both experiments (Fig. 4A & Dataset S1), further indicating a moderate level of similarities between transcriptional responses to MBX-4132 and knocking down tmRNA. Upregulated genes in both studies include aforementioned *whiB7* and *whiB6* (Dataset S1). Those encoding metal-sensing proteins (*cadI, cmtR, smtB,* and *mymT*) and efflux pumps (*ctpC, ctpG,* and *ctpJ*) were also induced in both experiments (Dataset S1). However, none of these regulons were found to be differentially expressed in either study. Nevertheless, using gene ontology (GO) analyses, we discovered that pathways associated with cadmium and copper responses were enriched in upregulated genes by both MBX-4132 and tmRNA KD (Fig. S5B&C). On the other hand, only one gene related to iron homeostasis, *mmpS5,* was found to be differentially expressed when tmRNA levels were reduced (Fig. 4E). We did not observe significantly altered expression of IdeR regulon genes either (Dataset S1). GO analyses revealed that four out of five most highly enriched pathways are related to iron homeostasis among genes induced by MBX-4132 treatment, whereas no such pathways were found to be significantly enriched by *ssr* KD (Fig. S5B&C). These observations indicate that some aspects of the metal homeostasis dysregulation seen after MBX-4132 treatment are likely due to inhibition of *trans*-translation, but dysregulation of the iron-responsive genes is caused by off-target effects.

### REFERENCES

1. Abdeldaim, G., Svensson, E., Blomberg, J., and Herrmann, B. (2016). Duplex detection of the Mycobacterium tuberculosis complex and medically important non-tuberculosis mycobacteria by real-time PCR based on the rnpB gene. APMIS *124*, 991–995. https://doi.org/10.1111/apm.12598.

2. Singh, A., Ubaid-ullah, S., Ramteke, A.K., and Batra, J.K. (2016). Influence of Conformation of M. tuberculosis RNase P Protein Subunit on Its Function. PLoS One *11*, e0153798. https://doi.org/10.1371/journal.pone.0153798.

3. Lin-Chao, S., Wei, C.-L., and Lin, Y.-T. (1999). RNase E is required for the maturation of ssrA RNA and normal ssrA RNA peptide-tagging activity. Proceedings of the National Academy of Sciences *96*, 12406–12411. https://doi.org/10.1073/pnas.96.22.12406.

4. Khare, G., Nangpal, P., and Tyagi, A.K. (2017). Differential Roles of Iron Storage Proteins in Maintaining the Iron Homeostasis in Mycobacterium tuberculosis. PLoS One *12*, e0169545. https://doi.org/10.1371/journal.pone.0169545.

5. Shyam, M., Shilkar, D., Verma, H., Dev, A., Sinha, B.N., Brucoli, F., Bhakta, S., and Jayaprakash, V. (2021). The Mycobactin Biosynthesis Pathway: A Prospective Therapeutic Target in the Battle against Tuberculosis. J. Med. Chem. *64*, 71–100. https://doi.org/10.1021/acs.jmedchem.0c01176.

6. Wells, R.M., Jones, C.M., Xi, Z., Speer, A., Danilchanka, O., Doornbos, K.S., Sun, P., Wu, F., Tian, C., and Niederweis, M. (2013). Discovery of a Siderophore Export System Essential for Virulence of Mycobacterium tuberculosis. PLOS Pathogens *9*, e1003120. https://doi.org/10.1371/journal.ppat.1003120.

7. Sritharan, M. (2016). Iron Homeostasis in Mycobacterium tuberculosis: Mechanistic Insights into Siderophore-Mediated Iron Uptake. Journal of Bacteriology *198*, 2399–2409. https://doi.org/10.1128/jb.00359-16.

8. Arnold, F.M., Weber, M.S., Gonda, I., Gallenito, M.J., Adenau, S., Egloff, P., Zimmermann, I., Hutter, C.A.J., Hürlimann, L.M., Peters, E.E., et al. (2020). The ABC exporter IrtAB imports and reduces mycobacterial siderophores. Nature *580*, 413–417. https://doi.org/10.1038/s41586-020-2136-9.

9. Rodriguez, G.M., Sharma, N., Biswas, A., and Sharma, N. (2022). The Iron Response of Mycobacterium tuberculosis and Its Implications for Tuberculosis Pathogenesis and Novel Therapeutics. Front. Cell. Infect. Microbiol. *12*. https://doi.org/10.3389/fcimb.2022.876667.

10. Rodriguez, G.M., Voskuil, M.I., Gold, B., Schoolnik, G.K., and Smith, I. (2002). ideR, an Essential Gene in Mycobacterium tuberculosis: Role of IdeR in Iron-Dependent Gene Expression, Iron Metabolism, and Oxidative Stress Response. Infect Immun *70*, 3371–3381. https://doi.org/10.1128/IAI.70.7.3371-3381.2002.

11. Shi, X., and Darwin, K.H. (2015). Copper homeostasis in Mycobacterium tuberculosis. Metallomics *7*, 929–934. https://doi.org/10.1039/c4mt00305e.

12. Shi, X., Festa, R.A., Ioerger, T.R., Butler-Wu, S., Sacchettini, J.C., Darwin, K.H., and Samanovic, M.I. (2014). The Copper-Responsive RicR Regulon Contributes to Mycobacterium tuberculosis Virulence. mBio *5*, e00876-13. https://doi.org/10.1128/mBio.00876-13.

13. Xu, J., Ma, S., Huang, Y., Zhang, Q., Huang, L., Xu, H., Suleiman, I.M., Li, P., Wang, Z., and Xie, J. (2024). Mycobacterium marinum MMAR_0267-regulated copper utilization facilitates bacterial escape from phagolysosome. Commun Biol *7*, 1–15. https://doi.org/10.1038/s42003-024-06860-9.

14. Wolschendorf, F., Ackart, D., Shrestha, T.B., Hascall-Dove, L., Nolan, S., Lamichhane, G., Wang, Y., Bossmann, S.H., Basaraba, R.J., and Niederweis, M. (2011).

Copper resistance is essential for virulence of Mycobacterium tuberculosis. Proceedings of the National Academy of Sciences *108*, 1621–1626. https://doi.org/10.1073/pnas.1009261108.

15. Goethe, E., Laarmann, K., Lührs, J., Jarek, M., Meens, J., Lewin, A., and Goethe, R. (2020). Critical Role of Zur and SmtB in Zinc Homeostasis of Mycobacterium smegmatis. mSystems *5*, e00880-19. https://doi.org/10.1128/mSystems.00880-19.

16. Maciąg, A., Dainese, E., Rodriguez, G.M., Milano, A., Provvedi, R., Pasca, M.R., Smith, I., Palù, G., Riccardi, G., and Manganelli, R. (2007). Global Analysis of the Mycobacterium tuberculosis Zur (FurB) Regulon. Journal of Bacteriology *189*, 730–740. https://doi.org/10.1128/jb.01190-06.

17. Wang, S., Fang, R., Wang, H., Li, X., Xing, J., Li, Z., and Song, N. (2024). The role of transcriptional regulators in metal ion homeostasis of Mycobacterium tuberculosis. Front Cell Infect Microbiol *14*, 1360880. https://doi.org/10.3389/fcimb.2024.1360880.

18. Darwin, K.H. (2015). *Mycobacterium tuberculosis* and Copper: A Newly Appreciated Defense against an Old Foe?*. Journal of Biological Chemistry *290*, 18962–18966. https://doi.org/10.1074/jbc.R115.640193.

19. Botella, H., Peyron, P., Levillain, F., Poincloux, R., Poquet, Y., Brandli, I., Wang, C., Tailleux, L., Tilleul, S., Charrière, G.M., et al. (2011). Mycobacterial P1-Type ATPases Mediate Resistance to Zinc Poisoning in Human Macrophages. Cell Host Microbe *10*, 248–259. https://doi.org/10.1016/j.chom.2011.08.006.

20. Raimunda, D., Long, J.E., Padilla-Benavides, T., Sassetti, C.M., and Argüello, J.M. (2014). Differential roles for the / transporting ATPases, CtpD and CtpJ, in ycobacterium tuberculosis virulence. Molecular Microbiology *91*, 185–197. https://doi.org/10.1111/mmi.12454.

21. Li, S., Poulton, N.C., Chang, J.S., Azadian, Z.A., DeJesus, M.A., Ruecker, N., Zimmerman, M.D., Eckartt, K.A., Bosch, B., Engelhart, C.A., et al. (2022). CRISPRi chemical genetics and comparative genomics identify genes mediating drug potency in Mycobacterium tuberculosis. Nat Microbiol *7*, 766–779. https://doi.org/10.1038/s41564-022-01130-y.

22. Rock, J.M., Hopkins, F.F., Chavez, A., Diallo, M., Chase, M.R., Gerrick, E.R., Pritchard, J.R., Church, G.M., Rubin, E.J., Sassetti, C.M., et al. (2017). Programmable transcriptional repression in mycobacteria using an orthogonal CRISPR interference platform. Nat Microbiol *2*, 1–9. https://doi.org/10.1038/nmicrobiol.2016.274.

23. Madsen, C.T., Jakobsen, L., Buriánková, K., Doucet-Populaire, F., Pernodet, J.-L., and Douthwaite, S. (2005). Methyltransferase Erm(37) Slips on rRNA to Confer Atypical Resistance in *Mycobacterium tuberculosis**. Journal of Biological Chemistry *280*, 38942–38947. https://doi.org/10.1074/jbc.M505727200.

24. Poulton, N.C., DeJesus, M.A., Munsamy-Govender, V., Kanai, M., Roberts, C.G., Azadian, Z.A., Bosch, B., Lin, K.M., Li, S., and Rock, J.M. (2024). Beyond antibiotic resistance: The *whiB7* transcription factor coordinates an adaptive response to alanine starvation in mycobacteria. Cell Chemical Biology *31*, 669-682.e7. https://doi.org/10.1016/j.chembiol.2023.12.020.

25. Rudra, P., Hurst-Hess, K.R., Cotten, K.L., Partida-Miranda, A., and Ghosh, P. (2020). Mycobacterial HflX is a ribosome splitting factor that mediates antibiotic resistance. Proceedings of the National Academy of Sciences *117*, 629–634. https://doi.org/10.1073/pnas.1906748117.

26. Jolliffe, I.T., and Cadima, J. (2016). Principal component analysis: a review and recent developments. Philos Trans A Math Phys Eng Sci *374*, 20150202. https://doi.org/10.1098/rsta.2015.0202.
